# Common and separable neural alterations in substance use disorders: evidence from coordinate-based meta-analyses of functional neuroimaging studies in human

**DOI:** 10.1101/2020.02.19.956755

**Authors:** Benjamin Klugah-Brown, Xin Di, Jana Zweerings, Klaus Mathiak, Benjamin Becker, Bharat Biswal

**Affiliations:** The Clinical Hospital of Chengdu Brain Science Institute, MOE Key Laboratory for Neuroinformation, Center for Information in Medicine, School of Life Science and Technology, University of Electronic Science and Technology of China, No.2006, Xiyuan Avenue, West Hi-Tech Zone, Chengdu, Sichuan 611731, China; Department of Biomedical Engineering, New Jersey Institute of Technology, 619 Fenster Hall, Newark, NJ 07102, USA; Department of Psychiatry, Psychotherapy and Psychosomatics, Faculty of Medicine, RWTH Aachen, Pauwelstrasse 30, 52074, Aachen, Germany; JARA Translational Brain Medicine, RWTH Aachen, Pauwelstrasse 30, 52074, Aachen, Germany

**Keywords:** substance use disorder, nicotine, alcohol, cocaine, cannabis, reward, cognition, striatum, frontal cortex, functional magnetic resonance imaging

## Abstract

Delineating common and separable neural alterations in substance use disorders (SUD) is imperative to understand the neurobiological basis of the addictive process and to inform substance-specific treatment strategies. Given numerous functional MRI (fMRI) studies in different SUDs, meta-analysis could provide an opportunity to determine robust shared and substance-specific alterations. The present study employed a coordinate-based meta-analysis covering fMRI studies in individuals with addictive cocaine, cannabis, alcohol, and nicotine use. The primary meta-analysis demonstrated common alterations in primary dorsal striatal, and frontal circuits engaged in reward/salience processing, habit formation, and executive control across different substances and task-paradigms. Subsequent sub-analyses revealed substance-specific alterations in frontal and limbic regions, with marked frontal and insula-thalamic alterations in alcohol and nicotine use disorders respectively. Finally, examining task-specific alterations across substances revealed pronounced frontal alterations during cognitive processes yet stronger striatal alterations during reward-related processes. Together the findings emphasize the role of dysregulations in striato-frontal circuits and dissociable contributions of these systems in the domains of reward-related and cognitive processes which may contribute to substance-specific behavioral alterations.

## 1 Introduction

Problematic use of illicit and licit drugs and substance use disorders represent a major challenge for society, in terms of individual suffering and socio-economic costs [1]–[3]. Substance use disorders are estimated to contribute to 20% to the world mental illness [4] and recent large scale surveys estimate that worldwide over 35 million people fulfill the criteria for a substance use disorder [5]. Disorders related to alcohol, nicotine, stimulant (e.g. cocaine), and cannabis use are among the most prevalent. Despite increasing treatment demand for problematic use of these substances [6] treatment options still remain limited and of moderate efficacy [7].

Based on animal models and human neuroimaging research substance use disorders, particularly addiction as a common pathological endpoint, has been reconceptualized as a chronic relapsing disorder of the brain that is characterized by a preoccupation with drug-seeking and taking, compulsive use, loss of behavioral control, and withdrawal (DSM-V). On the neural level, the transition from volitional use to problematic and ultimately compulsive use is driven by progressive dysregulations in the brain’s motivational and cognitive circuits, particularly the striato-frontal circuits engaged in incentive salience and reward processing, habit formation, and executive control [8]–[10].

Based on early animal studies demonstrating that the acute reinforcing effects of all drugs of potential abuse increase dopamine in the terminal regions of the mesocorticostriatal system including the ventral striatum [11] - which with repeated use may drive dysregulations in incentive salience and habit formation [9], [12] - most research emphasizes the common neuropathological endpoints across substances and substance use disorders. In line with animal models demonstrating that neuroplastic changes in the striatum mediate exaggerated salience to drug cues at the expense of natural rewards and habitual responses to cues repeatedly paired with the drug [13], exaggerated striatal drug cue reactivity and blunted striatal processing of non-drug rewards has been demonstrated in functional MRI studies in human drug users with regular and addictive use of different substances [14]–[18]. However, despite convergent evidence for striatal maladaptation across different substance use disorders, substance-specific predisposing factors [19]–[22] and addiction-related alterations have been increasingly recognized, such that frontal regions have been found to be differentially impacted by stimulant or opioid use [23] and neurocognitive deficits in domains associated with striato-frontal circuits such as inhibitory control and cognitive flexibility have been found to be differentially impacted by alcohol, stimulants, and cannabis [24], [25]. Further evidence for substance use disorder-specific brain alterations comes from a recent qualitative review suggesting that different addictions may be associated with alterations in distinct brain systems and particularly alterations in frontal regions appear to be substance-specific [26].

The differences might result from common versus substance-specific predisposing factors that render individuals vulnerable to develop escalation of use in general versus for a particular substance [27]. Furthermore, differences in the neurobiological effects of the substances may arise from the specific neurotoxic profiles and neurotransmitter systems. In addition, the substances engage different primary neurotransmitter systems [28], which is related to transmitter system-specific neuroadaptations for long term users. Also, the acute rewarding effects of all substances engage the dopamine system leading to the down-regulation of dopamine receptors [8] for chronic users. The acute effects of cannabis are mediated by the endocannabinoid system and regional-specific downregulation of the cannabinoid CB1 receptor [29], the acute effects of nicotine are primarily mediated by its stimulatory effects on neuronal nicotinic acetylcholine receptors (nAChRs) and long-term nicotine exposure leads to neuroplastic adaptations in nACh receptor expression [30], [31], and the acute effects of cocaine are primarily mediated by effects on the dopamine system and marked neuroplastic changes of striatal dopamine receptors have been consistently reported in cocaine use disorder [32], [33].

Despite emerging evidence for common but also substance-specific neurobiological alterations, most previous research emphasized common pathological pathways. Although the determination of common pathways of addiction may promote the development of general treatment approaches, the identification of substance-specific neurobiological mechanisms is essential to further enhance our understanding of predisposing factors and to develop specialized treatment options. To address common limitations of single studies such as low sample size, study-specific characteristics of the sample and inclusion of one substance only, the present study employed a meta-analytic approach covering previous task fMRI studies on alcohol, cannabis, cocaine, and nicotine substance use disorder to determine common and disorder-specific neural alterations. To this end, we conducted a quantitative coordinate-based meta-analysis (CBMA) covering previous fMRI studies in substance use populations employing whole-brain foci from the studies selected according to our inclusion criteria. The CBMA approach was preferred to other methods like image-based meta-analysis because it takes advantage of the published coordinates, and quantitatively provides a summary of the presented results under the specific research question, while the latter approach is limited by the availability of whole-brain images (most studies rarely share their statistical images). We first conducted a main ALE analysis to determine core regions that neurally underpin substance use disorders across substances. This was followed by sub-meta-analyses employing substance-specific subtraction and conjunction analysis to further specify common and substance-specific neural alterations as well as functional domain-specific alterations for reward and cognitive processes. Based on previous animal models and human imaging research, we hypothesized common alterations in striatal systems engaged in reward/motivation (ventral striatum) and habit formation (dorsal striatum) as well as partly dissociable effects on frontal systems engaged in executive control and behavioral regulation. Moreover, we examined whether the observed substance-specific alterations are driven by an interaction between the substance used and the class of task paradigms employed.

## 2 Methods

### 2.1 Literature selection

We obtained articles including four kinds of substances that are regularly abused namely cocaine, cannabis, alcohol, and nicotine (cigarettes or tobacco). Utilizing Scopus, PubMed, and Web of Science, peer-reviewed studies published between 1^st^ January 2000 and 1^st^ November 2019 were collected using the following search terms; “Alcohol” OR “Cocaine” OR “Cannabis” OR “Nicotine/Tobacco/cigarette “AND “Functional magnetic resonance imaging” OR “fMRI”. The reference list of the selected articles was inspected separately. We targeted articles that reported: whole-brain coordinates either in the main paper or supplementary material with stereoscope coordinates in either Talairach or MNI (Montreal Neurological Institute) space, comparisons between healthy controls and patients with substance dependency or heavy usage. The exclusion criteria were as follows: 1. Articles reporting only region-of-interest (ROI) results (if the study additionally reported whole-brain corrected findings these were included), 2. Articles with poly-drug users and high comorbidities with psychiatric or somatic disorders (e.g. schizophrenia or HIV), 3. Articles focusing on parental exposure, and 4. Articles reporting results from the exact same dataset from previous studies. The breakdown of article screening and exclusion for the main and sub-meta-analysis is shown in Figure 1.

**Figure 1.**
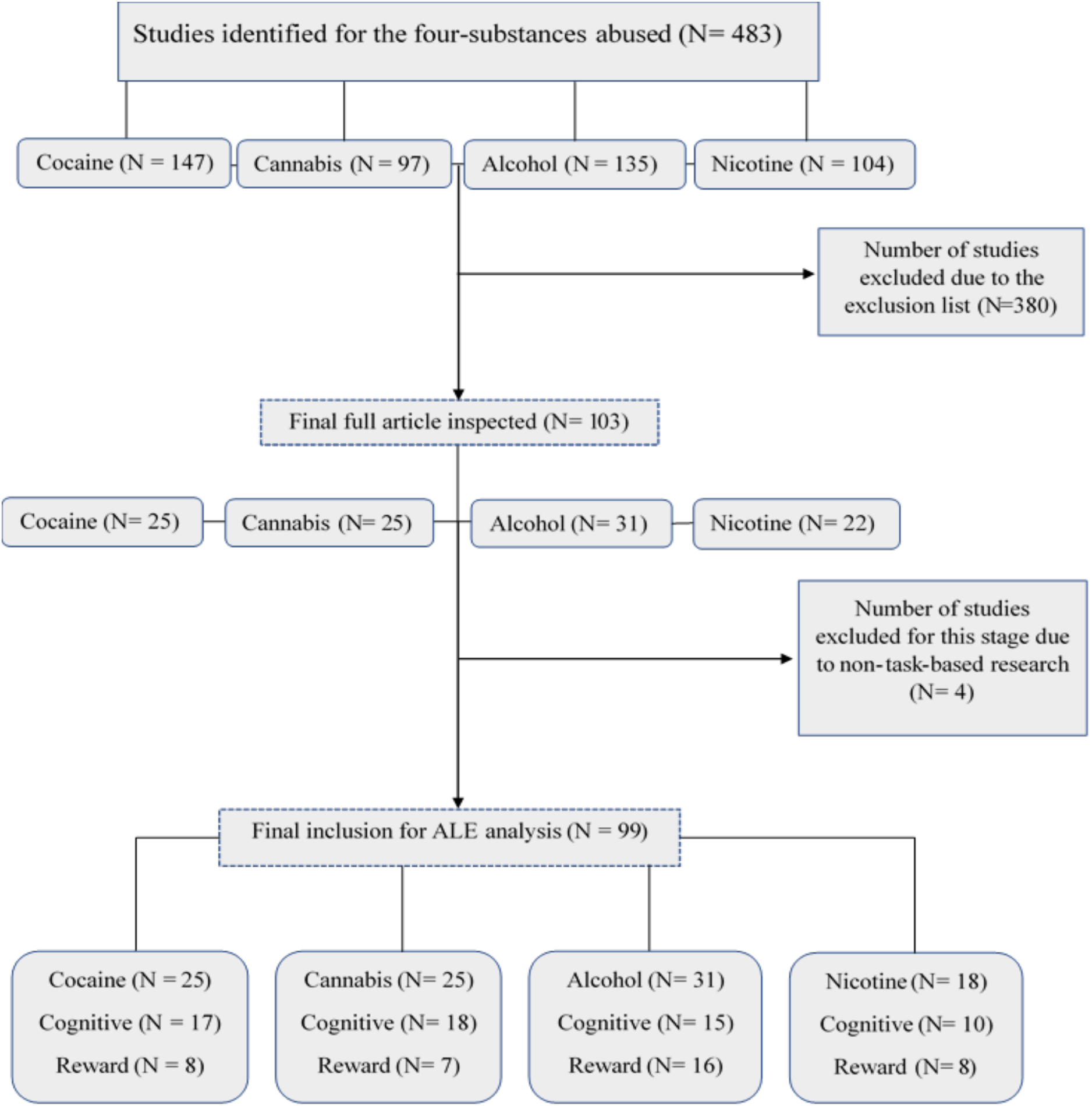
procedure for inclusion of articles

In total, 99 studies comprising 2763 reported foci were finally included in the meta-analysis. Articles included active as well as abstinent drug-using samples. To address the influence of age differences between the pairs of studies, we conducted a chi-square test using the Yates correction at p<0.05.

### 2.2 Study approach: Activation likelihood estimation (ALE)

Here, we used a two-step approach (1) to establish common brain regions that demonstrate brain functional alterations in substance use disorders, and (2) to investigate whether common regions are experimentally driven or solely by the effect of the drug in the studies.

We performed the coordinate-based analysis using the GingerALE 3.0.2 command-line version [34]–[36], to compute 1. Individual ALE for direct comparison 2. Subtraction and conjunction between pairs of studies. In each step of the analysis, all foci in MNI were converted to Talairach space. Also, to address the spatial uncertainty linked to the foci, the three-dimensional Gaussian probability distribution was applied to centers of the coordinates and images derived from the foci for all experiments obtained as modeled activation (MA). It is worth noting that ALE tries to find a strong agreement between the included experiments. This is achieved by computing the union between the generated maps while taking into account the disparities between true activation and noise. Cluster-level Family-wise error (cluster-FWE) [37] correction was applied to the ALE results to account for multiple comparisons. The ALE maps were initially thresholded using the uncorrected cluster forming p <0.001 and cluster-level threshold p<0.05.

#### Step 1: Direct comparison of ALEs in individual studies

We computed single group ALE analysis for the four substances, using the parameters described above. Coordinates from each substance group were combined as one study before computing the ALE. This was done to ascertain the overall maximum activations across the SUD.

#### Step 2: Subtraction and conjunction analysis between pairs of studies

We examined differences and overlaps between all six pairs of substances. The comparisons were done without specific *a priori* hypotheses (although co-use of the substances is often reported in research, (see [38])) in the following manner: cannabis versus cocaine, alcohol versus cocaine, alcohol versus cannabis, cannabis versus nicotine, cocaine versus nicotine, and nicotine versus alcohol. ALE analysis for individual substance groups was conducted first. The pooled foci were further used to compute the cluster-level FWE corrected maps and subsequently to obtain ALE images. Furthermore, we conducted a subtraction analysis to obtain the differences in ALE between the substance groups. In all of the results, clusters of brain regions were identified including the number of studies that contributed to them.

#### Step 3 Post-hoc analysis based on the functional domain

Based on the results in step 2, we divided pairs of studies that had conjunction (matched/overlapped pairs) activation into two categories of task paradigms that reflected distinct functional domains; that is, ‘Reward’ comprising the processing of reward, motivation, or anticipation and ‘Cognition’ with cognitive-control, behavior, and emotion tasks. This steps’ primary purpose was to investigate whether the conjunctions were influenced by the experimental design or solely by the substances abused. To this end we computed whole-brain ALE firstly for all studies based on the experimental task, secondly, we grouped the studies into patient and control studies and computed the cluster ALE to ascertain the group interactions due to the type of experimental paradigms. Using the chi-square test (Yates correction), we computed the statistical independence of tasks on the overlaps. The percentage contribution to each overlapped activation was also computed by taking the tally of each experimental paradigm in the pairs. The threshold for chi-square test was p<0.05.

## 3 Results

### 3.1 Included study sample characteristics

Out of the 99 studies included in the meta-analysis, cocaine studies contributed to 30% (828) of the total foci, while cannabis, alcohol, and nicotine contributed to 23.01% (637), 27.45% (760) and 19.44% (538) respectively of the total foci analyzed, indicating a bias towards a single substance due to an imbalanced data-base. Table 1 shows the demography of the four groups of substance abuse studies and the type of experiment or task used in each study group.

The combined studies yielded data from a total of 2777 substance users (mean (SD) age, 34.5 (12.05)) and 2643 control subjects (mean (SD) age, 31.63(11.63)), there was no significant difference between the age of the pairs of studies (cocaine versus cannabis *χ*^2^ = 0.0014, cocaine and alcohol *χ*^2^= 0.008, cocaine and nicotine *χ*^2^= 0.0205, cannabis, and alcohol, *χ*^2^= 0.0123, cannabis, and nicotine, *χ*^2^= 0.0005, alcohol, and nicotine, *χ*^2^= 0.0049).

**Table 1.** Subject characteristics for each study in a group

| Study source | Participants (N) |  | Age, Mean (SD) |  | Type of experiment/task |
| --- | --- | --- | --- | --- | --- |
|  | SUD | HC | SAB | HC |  |
| Cocaine studies |  |  |  |  |  |
| [39] (Barrós-Loscertales et al., 2011) | 16 | 16 | 34.38(7.15) | 34.2(8.86) | Stroop task |
| [40] (Barrós-Loscertales et al., 2019) | 30 | 28 | 35.9(6.31) | 38.89(10.5) | Stop-signal task |
| [41] (Bustamante et al., 2011) | 15 | 15 | 32.4(7.56) | 34.2(8.86) | verbal working memory task |
| [42] (Caldwell et al., 2015) | 219 | 87 | 34.9(8.08) | 32.15(9.07) | Moral judgement task |
| [43] (Crunelle et al., 2015) | 51 | 32 | 32(8) | 33(9) | Emotional face matching task |
| [44] (Ersche et al., 2013) | 18 | 18 | 34.3(7.2) | 32.7(6.9) | Stroop task |
| [45] (Garavan et al., 2000) | 31 | 17 | 34(0.5) | 26(0.7) | Working memory task |
| [46] (Ide et al., 2016) | 75 | 88 | 39.9(7.6) | 38.7(10.9) | Stop signal task |
| [47] (Kaag et al., 2016) | 40 | 51 | 31.3(7.9) | 31(8.5) | Fear conditioning paradigm |
| [48] (Kaag et al., 2018) | 59 | 58 | 31.4(7.6) | 30.5(8.1) | cue reactivity paradigm |
| [49] (Kirschner et al., 2018) | 22 | 28 | 29.73(7.99) | 28.2(6.72) | Prospective Imagery Task |
| [50] (Kober et al., 2016) | 30 | 73 | 43.78(13.06) | 32.22(11.06) | Craving/ emotional response task |
| [51] (Ma et al., 2015) | 13 | 10 | 37.4(5.3) | 35.2(7.3) | Go/NoGo task |
| [52] (McHugh et al., 2017) | 45 | 22 | 43.42(7.04) | 42.05(8.4) | Wisconsin Card Sorting Task |
| [53] (Marci et al., 1998) | 15 | 15 | 39(10.4) | 40.9(7.4) | Stroop task |
| [54] (F. G. Moeller et al., 2010) | 19 | 14 | 40.8(8.4) | 34.5(1.8) | Working memory task |
| [55] (S. J. Moeller et al., 2015) | 37 | 55 | 43.62(6.7) | 40.28(7.44) | Stroop task |
| [56] (S. J. Moeller, et al., 2015 (b)) | 33 | 20 | 43.55(8.3) | 39.6(5.5) | Inhibitory control task |
| [57] (S. J. Moeller et al., 2018) | 37 | 26 | 46.05(8.3) | 43.1(7.2) | Drug-Choice Task |
| [58] (Potenza et al., 2012) | 30 | 36 | 36.9(6.4) | 31.2(9) | Individualized scripts for stress |
| [59] (Sinha et al., 2005) | 20 | 8 | 38.75(4.77) | 32.8(4.74) | stress and neutral script |
| [60] (Tau et al., 2014) | 13 | 13 | 37.7(6.8) | 36.6(6) | Reward-based spatial learning task |
| [61] (Worhunsky et al., 2013) | 20 | 20 | 38.6(9.3) | 36.8(8.9) | Stroop task |
| [62] (Yip et al., 2016) | 20 | 21 | 38.6(9.29) | 34.57(11.99) | Monetary incentive delay task |
| [63] (Zhang et al., 2016) | 100 | 100 | 40.3(7.4) | 38(10.6) | Stop signal task. |
| Cannabis studies |  |  |  |  |  |
| [64] (Abdullaev et al., 2010) | 14 | 14 | 19.5(0.8) | 19.7(1.4) | Attention Network Task |
| [65] (Ames et al., 2013) | 16 | 17 | 21.15(1.9) | 20.27(2.3) | Implicit Association Test |
| [66] (Chang et al., 2006) | 24 | 19 | 28.77(2.81) | 30.57(1.83) | Non-verbal visual-attention task |
| [67] (Cousijn et al., 2014) | 32 | 41 | 21.65(2.4) | 22.25(2.35) | N-back task |
| [68] (De Bellis et al., 2013) | 15 | 41 | 16.4(7.3) | 16(1.2) | Decision-Reward Uncertainty task |
| [69] (Enzi et al., 2015) | 15 | 15 | 26.33(2.94) | 27.13(8.85) | Monetary Incentive Delay task |
| [70] (Filbey et al., 2016) | 53 | 68 | 30.66(7.48) | 31.41(10.2) | Cannabis cue-exposure task |
| [71] (Filbey et al., 2009) | 38 | 25 | 23.74(7.25) | 22.04(5.63) | Cue-elicited craving paradigm |
| [72] (Gilman et al., 2016) | 20 | 23 | 20.6(2.5) | 21.6(1.9) | Visual discrimination task |
| [73] (Gruber et al., 2009) | 15 | 15 | 25(8.8) | 26(9.0) | Masked affective tasks |
| [74] (Harding et al., 2012) | 21 | 21 | 36.5(8.8) | 31(11.7) | Multi-Source Interference Task |
| [75] (Heitzeg et al., 2015) | 20 | 20 | 19.84(1.45) | 20.51(1.26) | Emotion-arousal word task |
| [76] (Kanayama et al., 2004) | 12 | 10 | 37.9(7.4) | 27.8(7.9) | Working memory task |
| [77] (Kober et al, 2014) | 20 | 20 | 26.65(9.81) | 29.2(10.06) | Stroop task |
| [78] (Li et al., 2005) | 8 | 18 | 36(7.5) | 37.2(5.6) | Script-guided imagery paradigm |
| [79] (Lopez-Larson et al., 2012) | 24 | 24 | 18.2(0.7) | 18(1.9) | Finger-tapping task |
| [80] (Ma et al., 2018) | 23 | 23 | 28.2(3.5) | 28.7(3.7) | N-back working memory task |
| [81] (Nestor et al., 2008) | 49 | 52 | 23.35(0.95) | 23.05(0.85) | Face-name task |
| [82] (Schweinsburg et al., 2008) | 15 | 17 | 18.1(0.7) | 17.9(1.0) | Spatial working memory task |
| [83] (Tervo-Clemmens et al., 2018) | 22 | 63 | 14.12(0.33) | 14.21(0.37) | Spatial working memory task |
| [84](Tervo-Clemmens et al., 2018) | 14 | 15 | 28.16(0.69) | 28.16(0.71) | Visuospatial working memory task |
| [85] (van Hell et al., 2010) | 14 | 13 | 24.5(4.45) | 24(2.7) | Monetary reward task was |
| [86] (Wesley et al., 2011) | 16 | 16 | 26.4(3.6) | 26.6(6.1) | Iowa gambling task |
| [18] (Zimmermann et al., 2019) | 23 | 23 | 23.86(3.36) | 23.67(2.88) | Interpersonal Touch Paradigm |
| [87] (Zimmermann et al., 2017) | 23 | 20 | 21.24(2.59) | 21.1(3.61) | Event-related cognitive reappraisal |
| **Alcohol studies** |  |  |  |  |  |
| [88] (Akine et al., 2007) | 9 | 9 | 34.6(6.5) | 36.2(7.2) | Long-term memory retrieval task |
| [89] (Bagga et al., 2014) | 18 | 18 | 36.5(5.0) | 35.2(3.7) | Abstract reasoning task |
| [90] (Beylergil et al., 2017) | 34 | 26 | 44.73(8.3) | 41.92(9.6) | Reward-guided decision-making task |
| [91] (Brumback et al., 2015) | 22 | 16 | 17.93(0.7) | 17.42(0.7) | Alcohol pictures cue reactivity task |
| [92] (Chanraud et al., 2009) | 24 | 24 | 47.8(7.7) | 45(5.6) | Free and Cued Selective Reminding Test<br>verbal episodic memory assessment |
| [93] (Dager et al., 2014) | 23 | 33 | 18.9(0.6) | 18.7(0.4) | Figural memory task |
| [94] (Deserno et al., 2014) | 13 | 14 | 45.08(6) | 43.86(9.2) | Reversal learning task Reversal |
| [95] (Gilman et al., 2015) | 18 | 18 | 37.7(7.8) | 34.5(8.0) | Risk-taking task |
| [96] (Gorka et al, 2019) | 38 | 27 | 23.8(3.0) | 24.3(2.8) | Startle threat task |
| [97] (Grodin et al., 2019) | 24 | 22 | 36.41(14) | 32.29(9.9) | Cue reactivity task |
| [98] (Grodin et al., 2016) | 17 | 17 | 32.25(6.9) | 27.72(4.3) | Monetary incentive delay task |
| [99] (Grüsser et al., 2004) | 10 | 10 | 36(11.0) | 41(8.0) | Cue response task |
| [100] (Andreas Heinz et al., 2007) | 12 | 12 | 39(7.0) | 40(8.0) | Cue response task |
| [101] (Hermann et al., 2006) | 10 | 10 | 40(7.0) | 38(5.0) | Cue response task |
| [102] (Hu et al., 2015) | 24 | 70 | 38.7(8.3) | 35.1(9.9) | Stop signal task |
| [103] (Jang et al., 2007) | 20 | 20 | 43.5(6.0) | 44.5(7.4) | Mixed cognitive tests |
| [104] (Jansen et al., 2019) | 39 | 39 | 41.64(8.6) | 44.06(11.0) | Emotion reappraisal task |
| [105] (Kienast et al., 2013) | 11 | 13 | 41.9(7.0) | 43.2(9.5) | Emotional task |
| [106] (Maurage et al., 2013) | 12 | 12 | 24.2(4.5) | 23.4(4.2) | Two-alternative forced choice task |
| [107] (M.-S. Park et al., 2007) | 9 | 9 | 23.22(2.5) | 23(2.6) | Cue response task |
| [108] (Reiter et al., 2016) | 43 | 35 | 44.42 (10.21) | 42.00(10.49) | Anticorrelated decision-making task |
| [109] (Sjoerds et al., 2013) | 31 | 19 | 48.5(8.5) | 47.7(11.0) | Instrumental learning task was |
| [110] (Squeglia et al., 2011) | 40 | 55 | 17.9(0.9) | 17.88(1.0) | Spatial working memory task |
| [111] (Wesley et al., 2017) | 24 | 11 | 33.3(8.4) | 28.8(7.8) | The N-Back working memory task |
| [112] (Wetherill et al., 2013) | 40 | 20 | 18.4(2.1) | 18.3(1.4) | Go/No-go task |
| [113] (Wiers et al., 2015) | 38 | 17 | 44.39(7.3) | 42.71(9.2) | Cue reactivity task |
| [114] (Worbe et al., 2014) | 19 | 21 | 23.21(3.52) | 24.14(3.13) | Risk-taking task |
| [115] (J. Wrase et al., 2002) | 37 | 44 | 43.5(8.7) | 34.2(9.3) | Cue reactivity task |
| [116] (Jana Wrase et al., 2007) | 16 | 16 | 42.38(7.52) | 39.94(8.59) | Monetary incentive delay task |
| [117] (Yang et al., 2013) | 15 | 15 | 42.3(7.1) | 45.5(8.5) | Anticipatory anxiety paradigm |
| [118] (Yoon et al., 2009) | 12 | 12 | 32(5.2) | 31(6.2) | Memory encoding tasks |
| **Nicotine studies** |  |  |  |  |  |
| [119] (Artiges et al., 2009) | 13 | 13 | 26(4.0) | 24(4.0) | Smoking cues emotion recognition task |
| [120] (Carroll et al., 2015) | 23 | 19 | 35(10) | 30.2(7.2) | Speeded Flanker task |
| [121] (Bühler et al., 2010) | 21 | 21 | 28(4.3) | 25.7(6.1) | Event-related instrumental motivation task |
| [122] (Galván et al., 2013) | 18 | 25 | 19.47(1.33) | 19.08(1.15) | Balloon Analogue Risk Task |
| [123] (Hong et al., 2017) | 17 | 16 | 39.9(4.9) | 39.2(5.2) | Cue reactivity task |
| [124] (Kobiella et al., 2014) | 27 | 33 | 41.3(7.9) | 41.3(7.9) | Intertemporal choice task |
| [125] (Okuyemi et al., 2006) | 17 | 17 | 37.65(9.4) | 35.8(10.95) | Cue viewing task |
| [126] (Lawn et al., 2019) | 19 | 19 | 29.5(10.7) | 22.7(4.4) | Value-based decision-making task |
| [127] (Lesage et al., 2017) | 24 | 20 | 35.8(9.9) | 30.4(7.2) | Probabilistic reversal learning task |
| [128] (Lieberman et al., 2018) | 5 | 5 | 21.7(3.8) | 21.7(3.8) | cue viewing task |
| [129] (Luijten et al., 2013) | 25 | 23 | 22.56(2.84) | 21.74(1.82) | Go/NoGo task |
| [130] (Luo et al., 2011) | 35 | 36 | 34.1(7.9) | 31.3(7.1) | Adaptive intertemporal choice task |
| [131] (Maynard et al., 2017) | 39 | 19 | 21.95(3.5) | 24(3) | Memory task |
| [132] (Rose et al., 2013) | 28 | 28 | 32.68(10.02) | 30.11(7.83) | Monetary incentive delay task |
| [133] (Rubinstein et al., 2011) | 12 | 12 | 16(1.4) | 16(1.4) | Cue Reactivity Paradigm |
| [134] (Wagner et al., 2011) | 17 | 17 | 23.1(NA) | 21.4(NA) | Cue response task |
| [135] (Weywadt et al., 2017) | 81 | 38 | 59(1.5) | 61(1.36) | Go/No-Go task |
| [136] (Yalachkov et al., 2012) | 15 | 15 | 28.3(3.7) | 27(5.01) | Visual stimuli |

### 3.2 Combined and subtraction ALE analysis

In the combined analysis, we found that most of the activations were located in the dorsal striatum, limbic lobe, and the prefrontal cortex represented by the caudate and putamen, anterior cingulate cortex (ACC) and the inferior/superior/medial frontal gyrus (see Figure 2, detailed coordinates forming the maximum cluster peaks for combined ALE are listed in Table 2). In addition, the subtraction analysis yielded significant differences between pairs of studies. The alcohol group showed greater alterations in the left middle frontal gyrus compared to the cannabis group that was mostly characterized by alterations in the right caudate, right insula, right superior frontal gyrus, and right inferior frontal gyrus. Also, nicotine associated changes were greater in the left caudate and left anterior cingulate (see Figure 3). The detailed list of coordinates for the subtraction analysis is shown in Supplementary Table 1.

**Table 2.**
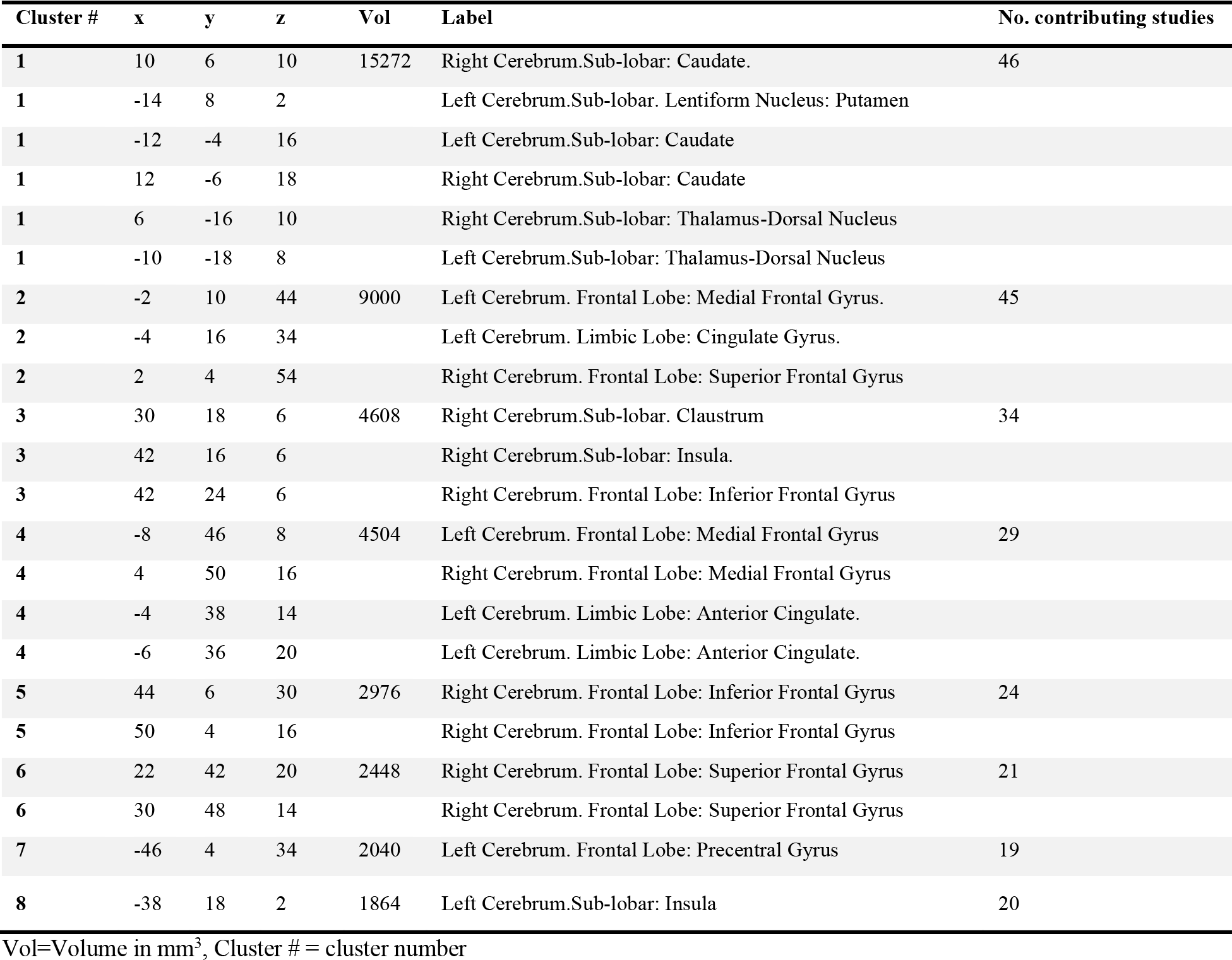
Detailed peak coordinates for the combined studies with the number of clusters for each volume

**Figure 2.**
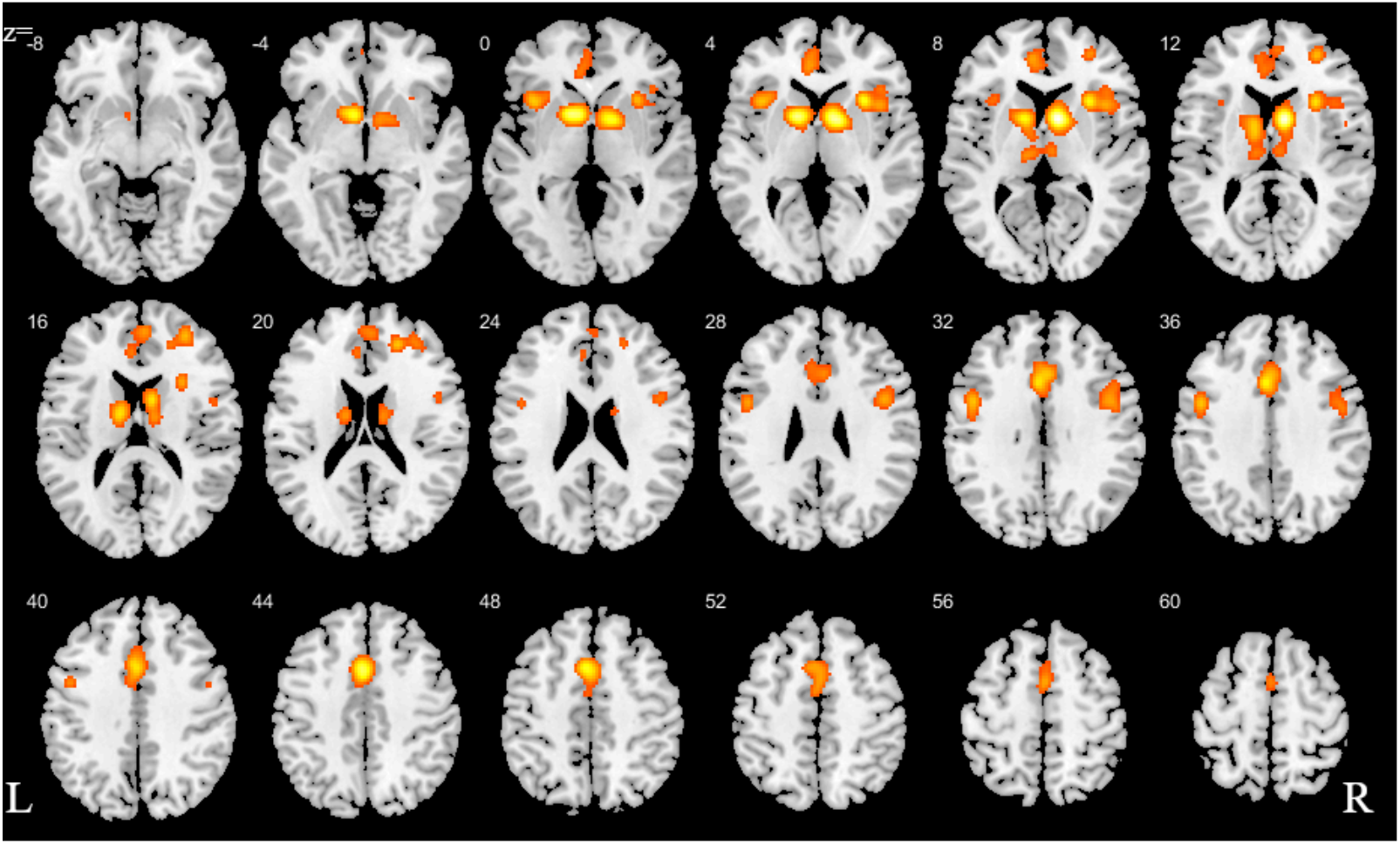
ALE for combined studies. All slices in transverse view with ascending slice number, uncorrected cluster forming p <0.001 and cluster-level threshold p<0.05.

**Figure 3.**
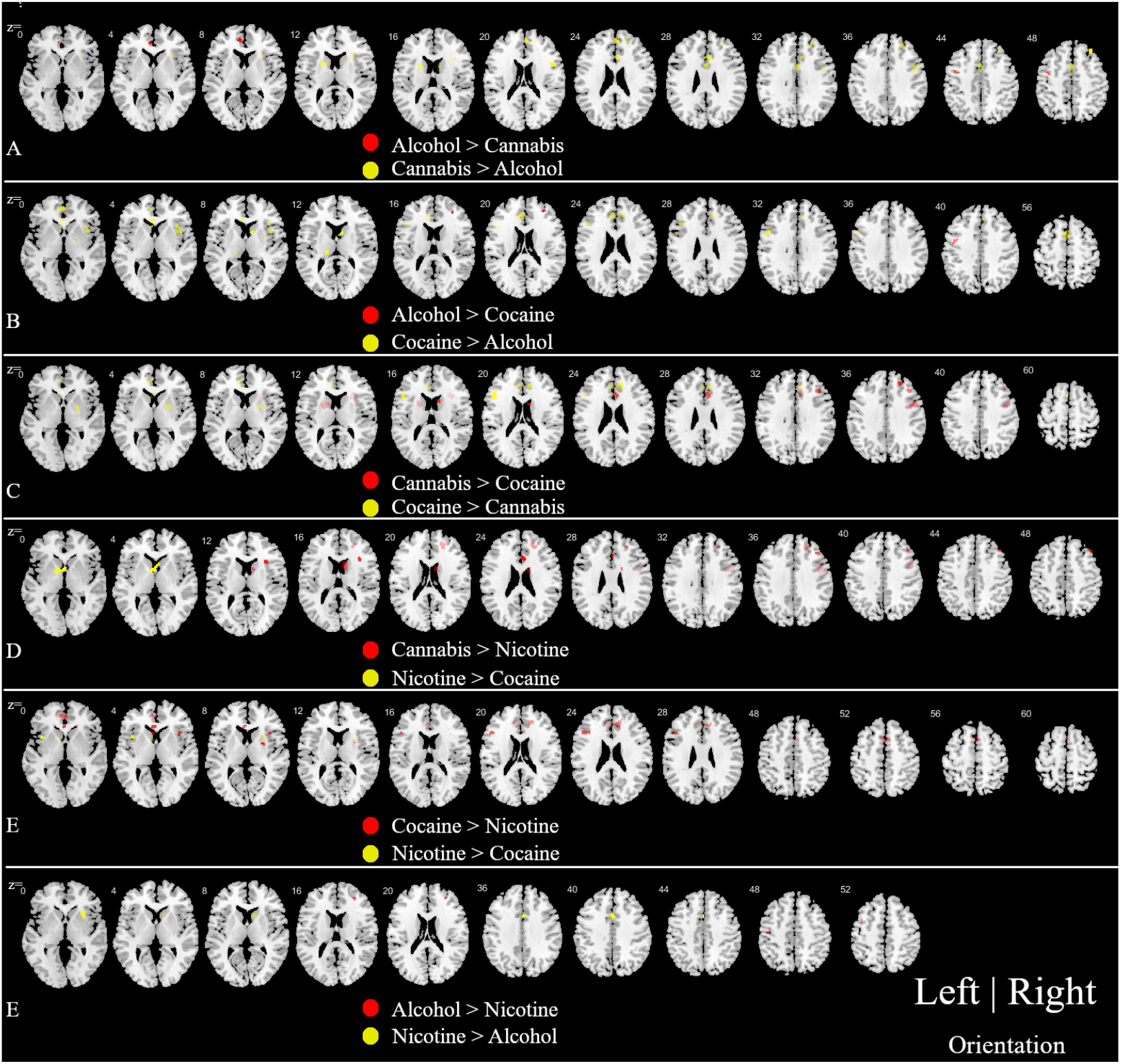
Subtraction analysis between pairs of studies.‘>’ symbol indicates where ALE peaks are greater in one study compared to the other. Cluster forming (p<0.001) and cluster-level threshold (p<0.05). Details on peak coordinates are shown in supplementary Table 1.

### 3.3 ALE results for conjugation between pairs of studies

We obtained the ALE images between each pair of studies as indicated as step 2 in the method section. In the alcohol versus the cocaine group, 8 clusters were obtained from the pooled foci results; we found two clusters - one in the frontal gyrus and the other in the dorsal striatum (Figure 4A). Similarly, the contrast between cannabis and cocaine revealed 4 significant clusters, exhibiting an overlap of one with the striatum (dorsal) and of two clusters with the frontal lobe (Superior Frontal Gyrus and Medial Frontal Gyrus; Figure 4B). We found no conjunction between alcohol and cannabis. The rest of the conjunctions are shown in Figure 4C-E with their details in Table 3.

**Table 3.** Peak ALE coordinates for the paired conjunction.

| Cluster # | x | y | z | Vol | Label | No. contributing studies |
| --- | --- | --- | --- | --- | --- | --- |
| Conjunction: Alcohol versus Cocaine |  |  |  |  |  |  |
| 1 | 14 | 6 | 4 | 592 | Right Cerebrum.Sub-lobar: Putamen | 3 |
| 1 | 8 | 4 | -2 |  | Right Cerebrum.Sub-lobar: Caudate |  |
| 2 | -8 | 48 | 8 | 496 | Left Cerebrum. Frontal Lobe: Medial Frontal Gyrus | 3 |
| 2 | -4 | 48 | 6 |  | Left Cerebrum. Limbic Lobe: Anterior Cingulate |  |
| 2 | -4 | 44 | 2 |  | Left Cerebrum. Limbic Lobe: Anterior Cingulate |  |
| 2 | -8 | 36 | 2 |  | Left Cerebrum. Limbic Lobe: Anterior Cingulate. |  |
| Conjunction: Cannabis versus Cocaine |  |  |  |  |  |  |
| 1 | 10 | 6 | 10 | 1000 | Right Cerebrum.Sub-lobar: Caudate | 8 |
| 2 | -2 | 12 | 50 | 968 | Left Cerebrum. Frontal Lobe: Superior Frontal Gyrus | 8 |
| 2 | -6 | 12 | 48 |  | Left Cerebrum. Frontal Lobe: Superior Frontal Gyrus |  |
| 3 | 20 | 40 | 20 | 208 | Right Cerebrum. Frontal Lobe: Medial Frontal Gyrus | 3 |
| 3 | 18 | 48 | 24 |  | Right Cerebrum. Frontal Lobe: Superior Frontal Gyrus |  |
| Conjunction: Cannabis versus Nicotine |  |  |  |  |  |  |
| 1 | 0 | 12 | 42 | 1104 | Left Cerebrum. Frontal Lobe: Medial Frontal Gyrus | 12 |
| 2 | 8 | 6 | 10 | 1024 | Right Cerebrum.Sub-lobar: Caudate | 7 |
| 3 | 30 | 18 | 8 | 1008 | Right Cerebrum.Sub-lobar: Claustrum | 3 |
| 4 | -12 | -4 | 18 | 984 | Left Cerebrum.Sub-lobar: Caudate | 7 |
| 4 | -12 | 6 | 12 |  | Left Cerebrum.Sub-lobar: Caudate |  |
| Conjunction: Cocaine versus Nicotine |  |  |  |  |  |  |
| 1 | 10 | 6 | 10 | 2088 | Right Cerebrum.Sub-lobar: Caudate | 9 |
| 1 | 20 | 4 | 0 |  | Right Cerebrum.Sub-lobar: Putamen |  |
| 2 | 36 | 16 | 4 | 184 | Right Cerebrum.Sub-lobar: Insula |  |
| Conjunction: Nicotine versus Alcohol |  |  |  |  |  |  |
| 1 | -14 | 10 | 0 | 1736 | Left Cerebrum.Sub-lobar: Putamen | 14 |
| 1 | -6 | 8 | -2 | 1080 | Left Cerebrum.Sub-lobar: Caudate | 7 |
| 2 | 10 | 8 | 2 |  | Right Cerebrum.Sub-lobar: Caudate |  |

Vol=Volume in mm<sup>3</sup>, Cluster # = cluster number, conjunction = overlap

**Figure 4.**
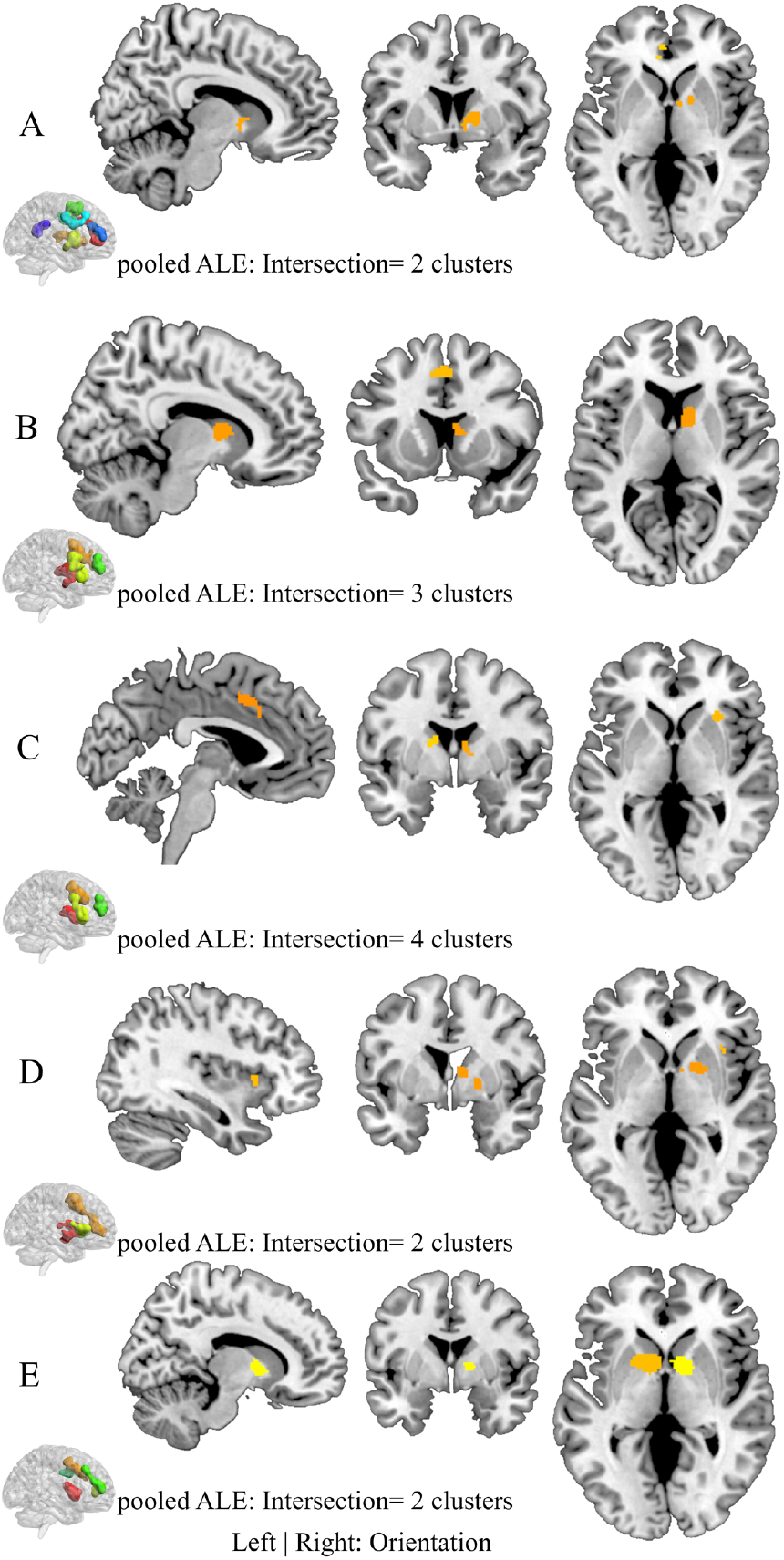
Conjunction analysis for each study pair. A) Alcohol versus Cocaine, B) Cannabis versus Cocaine, C) Cannabis versus Nicotine, D) Cocaine versus Nicotine and E) Nicotine versus Alcohol. All activations show the peak ALE p<0.001

### 3.4 Posthoc analysis: group-specific interaction

The group-specific interactions between SUD and controls are shown in Figures 5–7. This analysis revealed stronger activations in the frontal lobe during cognitive tasks for SUD. In addition, SUD exhibited stronger activations in the reward system in response to reward tasks, mostly in the dorsal striatum and the frontal lobes. Supplementary Table 4 shows the details of the coordinate information. Chi-square analysis results for overlaps based on tasks are shown in Supplementary Figure 3.

## 4 Discussion

The present meta-analytic approach employed a whole-brain coordinate-based meta-analysis to determine shared and substance-specific alterations in the most prevalent substance use disorders (alcohol, nicotine, cannabis, and cocaine) as determined in previous fMRI studies. Given that most previous studies could be classified in examining reward-related functions or cognitive/emotional functions, we additionally examined domain-specific alterations to determine whether dysregulations in distinct behavioral domains are neurally mediated by separable brain systems. In line with most overarching translational addiction models (e.g. [8]–[10]) the main meta-analysis demonstrated robust functional alterations in striato-frontal regions, particularly dorsal striatal regions involved in habit formation and compulsive behavior, as well as prefrontal regions including anterior cingulate, inferior frontal and medial prefrontal regions critically engaged in executive control and behavioral regulation. Exploratory substance-specific meta-analyses furthermore revealed a consistent pattern of altered neural processing in striatal and prefrontal regions for the separate substances, with some evidence for less frontal impairments in nicotine addiction. The comparative analyses between the substances moreover revealed some evidence for differential effects in fronto-striatal regions as well as limbic regions such as the ACC and the insular cortex. Further examining substance-specific differences revealed that the experimental paradigm was independent of the observed activations except for alcohol versus cocaine and nicotine versus alcohol, these were mostly driven by reward-based experiments. Cocaine and cannabis overlapped in the inferior frontal gyrus while alcohol-related patterns showed no overlap with cannabis for the contrast between reward-related and cognitive paradigms. Finally examining functional domain-specific alterations across all substances revealed that substance users demonstrated predominately striatal alterations during reward processes but frontal alterations during cognitive processes, suggesting that alterations in different behavioral domains are mediated by alterations in separable neural systems.

**Figure 5.**
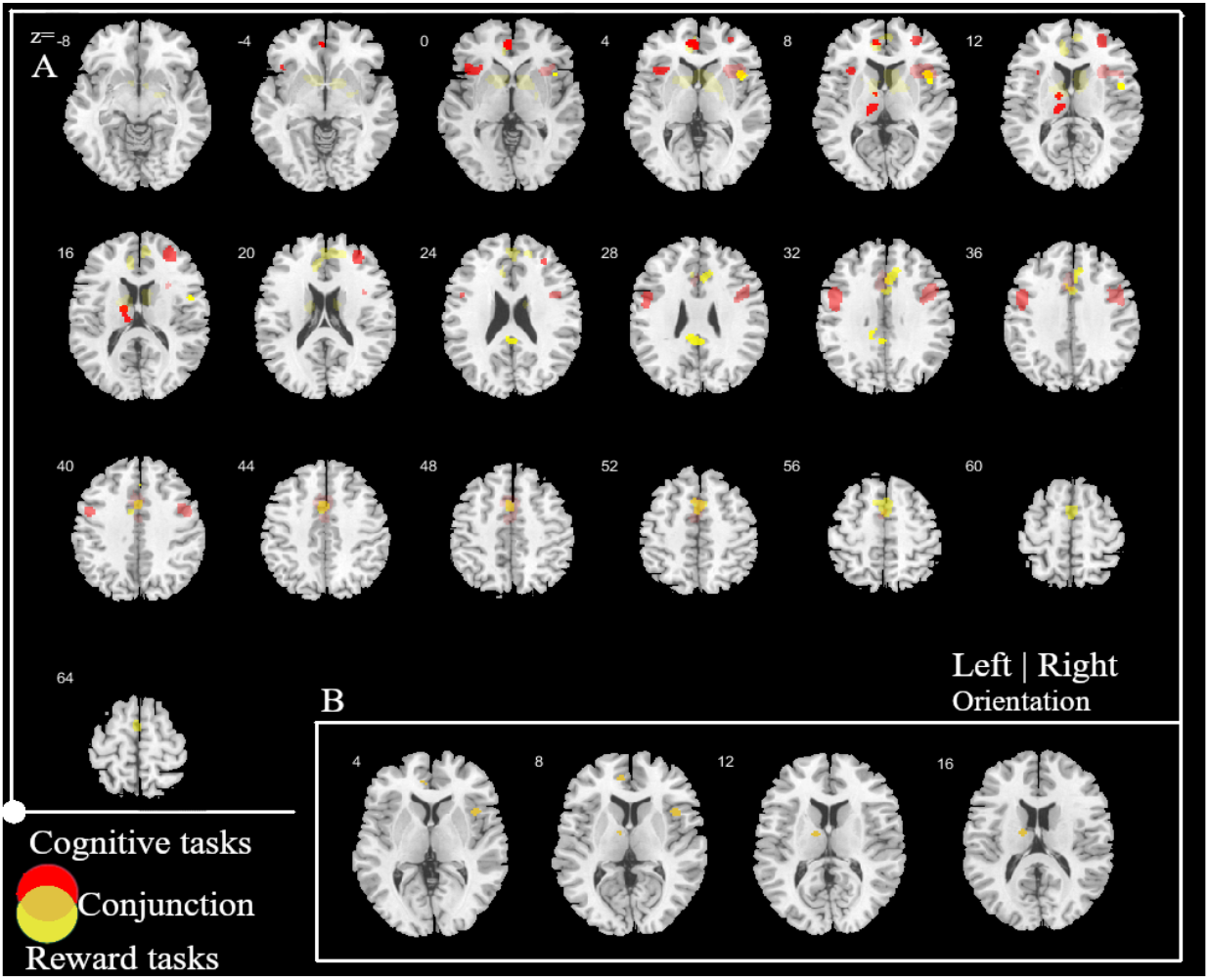
Conjunction analysis between cognitive-based tasks and reward-based tasks for the four substances. A) ALE peak alteration between the two types of tasks. Red (cognitive tasks > reward tasks); Yellow (reward tasks > cognitive tasks). B) conjunction between the two types of task paradigm

**Figure 6.**
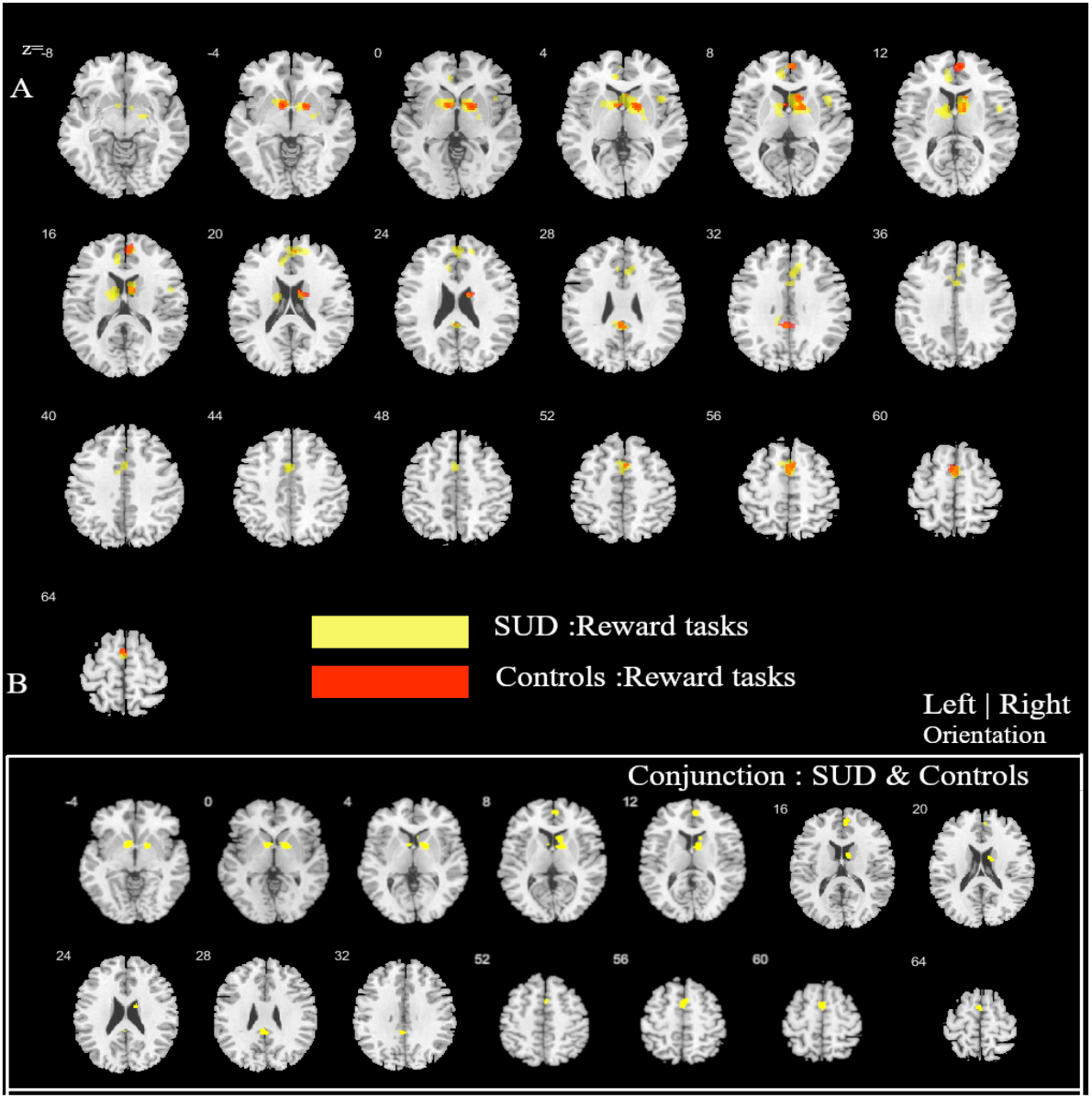
Conjunction analysis between individuals with substance use disorder (SUD) and controls in reward tasks for the four substances. A) ALE peak alteration between SUD and controls. Yellow (SUD > controls); Red (controls > SUD). B) shows the conjunction between the two groups.

**Figure 7.**
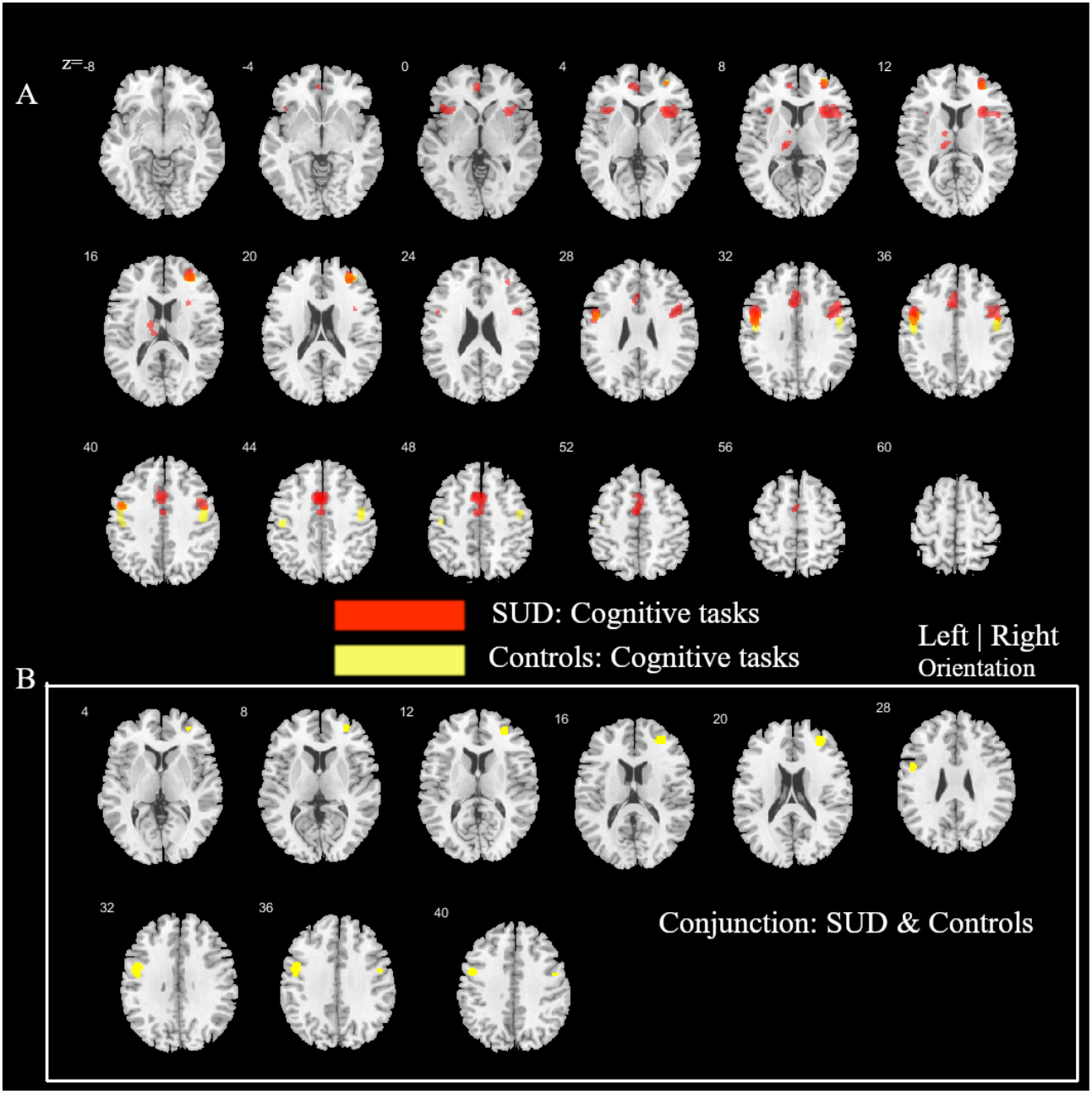
Conjunction analysis between individuals with substance use disorder (SUD) and controls in a cognitive task for the four substances. A) ALE peak alteration between SUD and controls. Yellow (SUD > controls); Red (controls > SUD). B) shows the conjunction between the two groups.

In general, findings from the present meta-analysis confirmed the extensive animal and human literature suggesting a critical role of the striato-frontal circuits in addiction. Neuroadaptations in this circuitry have been associated with behavioral dysregulations in the domains of incentive salience and reward processing, habit formation and executive control [8]–[10] and may underpin the progressive loss of control that represents a key symptom across substance use disorders. In line with the key symptomatic deficits in salience/reward processing and executive control deficits in substance use disorders most previous studies employed corresponding task-based paradigms examining associated neural processes. Comparing neural alterations in these domains across substances revealed stronger alterations in frontal regions during cognitive processes whereas alterations during reward/salience processing were neurally underpinned by stronger alterations in striatal regions and limbic regions, particularly the ACC. These findings resonate with the critical engagement of the frontal cortex in cognitive functions, including inhibitory control, decision making, and working memory which have been consistently found impaired in populations characterized by chronic substance use [137]–[139]. The ventral striatum represents one of the most commonly identified regions showing alterations in previous meta-analyses encompassing neuroimaging studies in addiction, specifically during reward-related processes [14], [17].

Together with the orbitofrontal cortex and the ACC the ventral striatum is engaged in evaluating the subjective value of stimuli in the environment [10] and has been associated with impulsive choices and trait impulsivity [140], [141]. Accordingly, alterations in this region may reflect adaptations in incentive-based learning processes that promote exaggerated salience attributed to the drug as well as deficits in controlling impulsive behavior. The dorsal striatum, on the other hand, has been strongly associated with habit learning and the transition from reward-driven to compulsive behavior in addiction [15], [16], [142] and may promote the development of compulsive drug use in the context of progressive loss of behavioral control [143]. Together, these findings emphasize that separable neural systems may mediate specific behavioral dysregulations and key diagnostic symptoms that characterize substance use disorders.

In line with the different neurobiological profiles of the substances and increasing evidence for substance-specific alterations, the present meta-analysis revealed evidence for differential alterations in the substance-using populations. Alcohol use disorders were characterized by stronger engagement of frontal regions compared to the other three substances examined which may point to differential neurocognitive deficits in substance use disorders with alcohol use disorder being characterized by marked impairments in the domains of cognitive flexibility and attention [24]. For nicotine use disorder stronger alterations in striatal and insular regions, yet comparably fewer alterations in frontal regions were observed. In addition, reduced self-control and the ability to obtain self-regulation are linked with SUDs such as alcohol intake, and other health-threatening behaviors, thus, stronger self-regulation moderates the usage of the substance [144], [145]. Alcohol, for instance, has been shown in studies to have a stronger effect in terms of self-regulation [146] which is the primary indicator of prefrontal processes. These findings may underscore the high addictive potential of tobacco, with rather moderate cognitive impairments in tobacco users [30] as well as an important role of the insula in nicotine addiction [147]. Given the high prevalence of nicotine addiction across populations with substance use disorders [148] these findings furthermore stress the importance to control for tobacco use in neuroimaging studies on addiction.

Moreover, overlapping alterations across the addictive substances in the dorsal striatum and the superior frontal gyrus were observed, suggesting that the recruitment of reward-based processes including anticipation, learning, cognitive control, and decision-making paths in contrast to the ventral striatum indicated in the reward and anticipation theories of addiction [149], [150]. Cannabis use was associated with greater alterations in the frontal region comparative to cocaine, suggesting that intoxication and decision making may be dominant in cannabis compared to cocaine use which may be predominately driven by dysregulated reward anticipation. Long term cannabis use has been associated with dysfunctional frontal processes related to cognition such as response time to decision cues and verbal memory [151], [152] while cocaine use has been repeatedly associated with marked dysregulations in motivation and executive functioning [153]–[155]. Moreover, alcohol and nicotine abuse shared common cortical regions and generally showed similar alterations to the other drugs, which may reflect the high rates of co-abuse of nicotine and alcohol in many drug abusers [156], [157]. However, alcohol use was additionally characterized by greater alterations across limbic areas including the ACC suggesting stronger dysregulations in salience processing, reinforcement learning and decision making in contrast to nicotine which may predominately disrupt striatal reward-related processes. Furthermore, as mentioned in the method of the analysis, we hypothesis that nicotine is a co-abused substance in all the psychoactive substances used [156], and nicotine would share common functional properties with the other substances. Remarkably, our results were consistent with the hypothesis but we additionally observed nicotine-specific alterations in the thalamus that were not observed in the main meta-analysis encompassing all drug classes. The thalamus exhibits particularly high expressions of nicotine-sensitive receptors in the brain, which may partly contribute to the nicotine-specific alterations observed in this region [158], [159]. This nicotine-specific finding may reflect that in addition to striatal reward-related dysregulations nicotine use induced neuroadaptations in the thalamic circuit play an important role and may promote response inhibition impairments observed in both, animal and human models of nicotine addiction [160]. The thalamus may, therefore, specifically contribute to nicotine abuse.

Although this meta-analysis revealed task-specific alterations between the various groups of studies, we could not conduct substance-specific sub-analyses comparing the different tasks within each substance group. This was due to the limited number of substance-specific studies. The within-group task-based analysis would have thrown more light on the substance-specific alteration for the individual substance use disorder studies. Furthermore, the reproducibility of common models for substance abuse in this meta-analysis may be also related to selection and publication bias of studies.

## 5 Conclusion

Summarizing, our analysis revealed consistent results with convergent evidence from animal and human studies demonstrating that addiction is characterized in neural systems subserving salience/reward processes, habit learning and executive control, including decision making and response inhibition, specifically the dorsal striatum and the prefrontal cortex. On the other hand, the present meta-analytic approach allowed us to determine substance-specific alterations in frontal, limbic as well as insular regions, pointing to specific pathological alterations in addition to shared pathological pathways.

## Acknowledgments

This work was supported by the National Key Research and Development Program of China (Grant No. 2018YFA0701400), National Natural Science Foundation of China (NSFC, No 91632117), and Science, Innovation and Technology Department of the Sichuan Province (2018JY0001).

