## Supplemental Material for "Common and separable neural alterations in substance use disorders: evidence from coordinate-based meta-analyses of functional neuroimaging studies in human"

#### Supplementary material

Benjamin Klugah-Brown[a], Xin Di[b], Jana Zweerings[c,d], Klaus Mathiak[c,d], Benjamin Becker[a*], Bharat Biswal[a,b*]

[a]The Clinical Hospital of Chengdu Brain Science Institute, MOE Key Laboratory for Neuroinformation, Center for Information in Medicine, School of Life Science and Technology, University of Electronic Science and Technology of China, No.2006, Xiyuan Avenue, West Hi-Tech Zone, Chengdu, Sichuan 611731, China

[b]Department of Biomedical Engineering, New Jersey Institute of Technology, 619 Fenster Hall, Newark, NJ 07102, USA

[c]Department of Psychiatry, Psychotherapy and Psychosomatics, Faculty of Medicine, RWTH Aachen, Pauwelstrasse 30, 52074, Aachen, Germany

[d]JARA Translational Brain Medicine, RWTH Aachen, Pauwelstrasse 30, 52074, Aachen, Germany

Corresponding authors
Bharat Biswal*


Benjamin Becker*


### Supplementary material

**Supplementary Table 1** Detailed peak coordinates from each subtraction in the pairs ">" signifies greater than between the pairs

| Cluster # | | | | | | |
|---|---|---|---|---|---|---|
| **Alcohol > Cannabis** | x | y | z | P | Z | Label |
| **1** | -40 | -2 | 46 | 0.01 | 2.326348 | Left Cerebrum. Frontal Lobe. Middle Frontal Gyrus |
| **2** | -8 | 46 | 6 | 0.013 | 2.226212 | Left Cerebrum. Frontal Lobe. Medial Frontal Gyrus. |
| **Cannabis > Alcohol** | | | | | | |
| **1** | 12.7 | 15 | 15.5 | 0 | 3.290527 | Right Cerebrum.Sub-lobar. Caudate |
| **1** | 12 | 6 | 18 | 0.002 | 2.878162 | Right Cerebrum.Sub-lobar. Caudate |
| **1** | 10 | 9 | 10 | 0.003 | 2.747781 | Right Cerebrum.Sub-lobar. Caudate. |
| **2** | 28 | 24 | 8 | 0.004 | 2.65207 | Right Cerebrum.Sub-lobar. Claustrum. |
| **2** | 30.3 | 24.3 | 6.5 | 0.002 | 2.878162 | Right Cerebrum.Sub-lobar. Insula. |
| **2** | 34 | 22.7 | 16.7 | 0.005 | 2.575829 | Right Cerebrum.Sub-lobar. Insula. |
| **2** | 34 | 18 | 18 | 0.006 | 2.512144 | Right Cerebrum.Sub-lobar. Insula. |
| **3** | 25.2 | 44.4 | 38 | 0 | 3.290527 | Right Cerebrum. Frontal Lobe. Superior Frontal Gyrus. |
| **3** | 23.2 | 46.3 | 32.7 | 0.002 | 2.878162 | Right Cerebrum. Frontal Lobe. Superior Frontal Gyrus. |
| **3** | 20.7 | 39.8 | 35.1 | 0.003 | 2.747781 | Right Cerebrum. Frontal Lobe. Superior Frontal Gyrus. |
| **3** | 14 | 52 | 22 | 0.009 | 2.365618 | Right Cerebrum. Frontal Lobe. Superior Frontal Gyrus. |
| **3** | 20 | 51 | 20 | 0.024 | 1.977369 | Right Cerebrum. Frontal Lobe. Superior Frontal Gyrus. |
| **4** | 46 | 4 | 36 | 0 | 3.290527 | Right Cerebrum. Frontal Lobe. Precentral Gyrus. |
| **4** | 46 | 10 | 38 | 0.002 | 2.878162 | Right Cerebrum. Frontal Lobe. Middle Frontal Gyrus. |
| **5** | 0 | 9 | 50 | 0.004 | 2.65207 | Left Cerebrum. Frontal Lobe. Superior Frontal Gyrus. |
| **6** | -18 | 6.7 | 15.3 | 0.007 | 2.457264 | Left Cerebrum.Sub-lobar. Caudate. |
| **6** | -22 | 12 | 16 | 0.008 | 2.408916 | Left Cerebrum.Sub-lobar. Claustrum. |
| **6** | -22.7 | 8 | 15.3 | 0.009 | 2.365618 | Left Cerebrum.Sub-lobar. Putamen |
| **7** | 8 | 22 | 24 | 0 | 3.290527 | Right Cerebrum. Limbic Lobe. Anterior Cingulate. |
| **8** | 2 | 5 | 33 | 0.005 | 2.575829 | Right Cerebrum. Limbic Lobe. Cingulate Gyrus. |
| **9** | 6 | 52 | 24 | 0.001 | 3.090232 | Right Cerebrum. Frontal Lobe. Superior Frontal Gyrus. |
| **10** | 44 | 12 | 22 | 0.024 | 1.977369 | Right Cerebrum. Frontal Lobe. Inferior Frontal Gyrus. |
| **10** | 52 | 10 | 18 | 0.036 | 1.799118 | Right Cerebrum. Frontal Lobe. Inferior Frontal Gyrus. |
| **10** | 52 | 6 | 22 | 0.042 | 1.727934 | Right Cerebrum. Frontal Lobe. Inferior Frontal Gyrus. |
| **11** | 32 | 38 | 50 | 0.001 | 3.090232 | Right Cerebrum. Frontal Lobe. Middle Frontal Gyrus. |
| **11** | 28 | 36 | 48 | 0.011 | 2.290368 | Right Cerebrum. Frontal Lobe. Middle Frontal Gyrus. |
| **11** | 32 | 34 | 50 | 0.012 | 2.257129 | Right Cerebrum. Frontal Lobe. Superior Frontal Gyrus. |
| **Alcohol > Cocaine** | | | | | | |
| **1** | -40 | 0 | 40 | 0.029 | 1.895698 | Left Cerebrum. Frontal Lobe. Precentral Gyrus. |
| **2** | 30 | 45 | 19 | 0.03 | 1.880794 | Right Cerebrum. Frontal Lobe. Middle Frontal Gyrus. |

| | | | | | | |
|---|---|---|---|---|---|---|
| **Cocaine > Alcohol** | | | | | | |
| **1** | 30.8 | -3.5 | -1.5 | 0 | 3.290527 | Right Cerebrum.Sub-lobar. Putamen |
| **1** | 19.6 | -10.8 | -11.6 | 0.003 | 2.747781 | Right Cerebrum. Limbic Lobe. Parahippocampal Gyrus. |
| **2** | -45.2 | 25 | 23.3 | 0.001 | 3.090232 | Left Cerebrum. Frontal Lobe. Middle Frontal Gyrus. |
| **2** | -46.7 | 25.3 | 18 | 0.001 | 3.090232 | Left Cerebrum. Frontal Lobe. Inferior Frontal Gyrus. |
| **2** | -36 | 20 | 28 | 0.018 | 2.096927 | Left Cerebrum. Frontal Lobe. Middle Frontal Gyrus. |
| **3** | 6 | 35 | 39 | 0.002 | 2.878162 | Right Cerebrum. Frontal Lobe. Medial Frontal Gyrus. |
| **3** | 8.7 | 38 | 36.7 | 1 | 0 | Right Cerebrum. Frontal Lobe. Medial Frontal Gyrus. |
| **3** | 14 | 38 | 26 | 0.006 | 2.512144 | Right Cerebrum. Frontal Lobe. Medial Frontal Gyrus. |
| **3** | 0 | 34 | 42 | 0.007 | 2.457264 | Left Cerebrum. Frontal Lobe. Superior Frontal Gyrus. |
| **3** | 8 | 38 | 24 | 0.011 | 2.290368 | Right Cerebrum. Limbic Lobe. Anterior Cingulate. |
| **4** | -11.2 | 38.4 | 20 | 0 | 3.290527 | Left Cerebrum. Frontal Lobe. Medial Frontal Gyrus. |
| **4** | -4.5 | 37.5 | 20.5 | 0.001 | 3.090232 | Left Cerebrum. Limbic Lobe. Anterior Cingulate. |
| **5** | -1.5 | 11 | 55 | 0 | 3.290527 | Left Cerebrum. Frontal Lobe. Superior Frontal Gyrus. |
| **5** | -2.7 | 16 | 57.3 | 0.001 | 3.090232 | Left Cerebrum. Frontal Lobe. Superior Frontal Gyrus. |
| **6** | 40 | 12 | 1 | 0.004 | 2.65207 | Right Cerebrum.Sub-lobar. Insula. |
| **6** | 39 | 17 | 3 | 0.01 | 2.326348 | Right Cerebrum.Sub-lobar. Insula. |
| **6** | 40 | 28 | 8 | 0.008 | 2.408916 | Right Cerebrum. Frontal Lobe. Inferior Frontal Gyrus. |
| **7** | -5 | 50 | -2 | 0.008 | 2.408916 | Left Cerebrum. Limbic Lobe. Anterior Cingulate. |
| **8** | -4 | 30 | 8 | 0.004 | 2.65207 | Left Cerebrum. Limbic Lobe. Anterior Cingulate. |
| **8** | -2 | 28 | 4 | 0.007 | 2.457264 | Left Cerebrum. Limbic Lobe. Anterior Cingulate. |
| **8** | -8 | 20 | 2 | 0.008 | 2.408916 | Left Cerebrum.Sub-lobar. Caudate |
| **8** | -4 | 24 | 2 | 0.01 | 2.326348 | Left Cerebrum.Sub-lobar. Caudate |
| **9** | -48 | 12 | 32 | 0.011 | 2.290368 | Left Cerebrum. Frontal Lobe. Middle Frontal Gyrus. |
| **10** | 12 | 12 | 10 | 0.024 | 1.977369 | Right Cerebrum.Sub-lobar. Caudate. |
| **10** | 8 | 8 | 12 | 0.025 | 1.959964 | Right Cerebrum.Sub-lobar. Caudate. |
| **11** | -16 | -24 | 10 | 0.019 | 2.074855 | Left Cerebrum.Sub-lobar. Thalamus. |
| **Cannabis > Cocaine** | | | | | | |
| **1** | 25 | 20 | 8 | 0 | 3.290527 | Right Cerebrum.Sub-lobar. Claustrum. |
| **1** | 26 | 20 | 2 | 0.007 | 2.457264 | Right Cerebrum.Sub-lobar. Claustrum. |
| **2** | -22 | 2 | 12 | 0.008 | 2.408916 | Left Cerebrum.Sub-lobar. Putamen |
| **2** | -16.7 | 5.3 | 15.3 | 0.011 | 2.290368 | Left Cerebrum.Sub-lobar. Caudate. |
| **3** | 48 | 8 | 36 | 0.007 | 2.457264 | Right Cerebrum. Frontal Lobe. Middle Frontal Gyrus. |
| **3** | 42 | 2 | 38 | 0.011 | 2.290368 | Right Cerebrum. Frontal Lobe. Middle Frontal Gyrus. |
| **4** | 42 | 30 | 42 | 0.003 | 2.747781 | Right Cerebrum. Frontal Lobe. Middle Frontal Gyrus. |
| **4** | 40 | 32 | 46 | 0.005 | 2.575829 | Right Cerebrum. Frontal Lobe. Middle Frontal Gyrus. |
| **4** | 44 | 30 | 46 | 0.008 | 2.408916 | Right Cerebrum. Frontal Lobe. Middle Frontal Gyrus. |
| **4** | 36 | 30 | 32 | 0.011 | 2.290368 | Right Cerebrum. Frontal Lobe. Middle Frontal Gyrus. |
| **5** | 6 | 20 | 24 | 0.003 | 2.747781 | Right Cerebrum. Limbic Lobe. Anterior Cingulate. |
| **6** | 14 | 10 | 18 | 0.004 | 2.65207 | Right Cerebrum.Sub-lobar. Caudate. |
| **7** | 22 | 44 | 38 | 0.014 | 2.197286 | Right Cerebrum. Frontal Lobe. Superior Frontal Gyrus. |

| | | | | | | |
|---|---|---|---|---|---|---|
| **7** | 24 | 40 | 38 | 0.015 | 2.17009 | Right Cerebrum. Frontal Lobe. Superior Frontal Gyrus. |
| **8** | -2 | 10 | 44 | 0.015 | 2.17009 | Left Cerebrum. Frontal Lobe. Medial Frontal Gyrus. |
| **Cocaine > Cannabis** | | | | | | |
| **1** | -4.8 | 35.6 | 21.1 | 0 | 3.290527 | Left Cerebrum. Limbic Lobe. Anterior Cingulate. |
| **1** | -9.4 | 39.5 | 12.2 | 0.003 | 2.747781 | Left Cerebrum. Limbic Lobe. Anterior Cingulate. |
| **1** | -8.7 | 34.7 | 4.7 | 0.141 | 0 | Left Cerebrum. Limbic Lobe. Anterior Cingulate. |
| **1** | -6 | 46 | -6 | 0.021 | 2.03352 | Left Cerebrum. Limbic Lobe. Anterior Cingulate. |
| **2** | 24 | 2 | 4 | 0.002 | 2.878162 | Right Cerebrum.Sub-lobar. Putamen |
| **2** | 24 | -8 | -10 | 0.007 | 2.457264 | Right Cerebrum.Sub-lobar. Amygdala |
| **2** | 26 | -6 | -2 | 0.023 | 1.995393 | Right Cerebrum.Sub-lobar. Putamen |
| **3** | 9.7 | 36 | 22.7 | 0.003 | 2.747781 | Right Cerebrum. Limbic Lobe. Anterior Cingulate. |
| **4** | -50 | 24 | 20 | 0.005 | 2.575829 | Left Cerebrum. Frontal Lobe. Inferior Frontal Gyrus. |
| **4** | -49.4 | 18 | 17.7 | 0.006 | 2.512144 | Left Cerebrum. Frontal Lobe. Inferior Frontal Gyrus. |
| **5** | -8 | 10 | 56 | 0.01 | 2.326348 | Left Cerebrum. Frontal Lobe. Superior Frontal Gyrus. |
| **5** | -5 | 15 | 57 | 0.02 | 2.053749 | Left Cerebrum. Frontal Lobe. Superior Frontal Gyrus. |
| **Cannabis > Nicotine** | | | | | | |
| **1** | 44.1 | 4.1 | 36.1 | 0 | 3.290527 | Right Cerebrum. Frontal Lobe. Precentral Gyrus. |
| **1** | 50 | 0 | 31 | 0.001 | 3.090232 | Right Cerebrum. Frontal Lobe. Precentral Gyrus. |
| **2** | 22 | 42 | 20 | 0.006 | 2.512144 | Right Cerebrum. Frontal Lobe. Superior Frontal Gyrus. |
| **2** | 24 | 40 | 16 | 0.008 | 2.408916 | Right Cerebrum. Frontal Lobe. Medial Frontal Gyrus. |
| **2** | 20 | 44 | 38 | 0.011 | 2.290368 | Right Cerebrum. Frontal Lobe. Superior Frontal Gyrus. |
| **2** | 24 | 38 | 34 | 0.014 | 2.197286 | Right Cerebrum. Frontal Lobe. Middle Frontal Gyrus. |
| **2** | 24 | 40 | 30 | 0.018 | 2.096927 | Right Cerebrum. Frontal Lobe. Superior Frontal Gyrus. |
| **3** | 10 | 8 | 20.7 | 0.004 | 2.65207 | Right Cerebrum.Sub-lobar. Caudate. |
| **3** | 12 | 13 | 18 | 0.003 | 2.747781 | Right Cerebrum.Sub-lobar. Caudate. |
| **3** | 18 | 2 | 26 | 0.008 | 2.408916 | Right Cerebrum.Sub-lobar. Caudate. |
| **4** | 41 | 30.5 | 46.5 | 0.019 | 2.074855 | Right Cerebrum. Frontal Lobe. Middle Frontal Gyrus. |
| **4** | 42 | 32 | 38 | 0.014 | 2.197286 | Right Cerebrum. Frontal Lobe. Middle Frontal Gyrus. |
| **5** | 36 | 20 | 14 | 0.005 | 2.575829 | Right Cerebrum.Sub-lobar. Insula. |
| **6** | 6 | 18 | 24 | 0 | 3.290527 | Right Cerebrum. Limbic Lobe. Anterior Cingulate. |
| **6** | 2 | 14 | 26 | 0.039 | 1.76241 | Right Cerebrum. Limbic Lobe. Cingulate Gyrus. |
| **Nicotine > Cannabis** | | | | | | |
| **1** | -2 | 6 | 2 | 0.004 | 2.65207 | Left Cerebrum.Sub-lobar. Caudate. |
| **1** | 0 | 6 | -2 | 0.005 | 2.575829 | Left Cerebrum. Limbic Lobe. Anterior Cingulate. |
| **1** | -8 | 4 | -2 | 0.008 | 2.408916 | Left Cerebrum.Sub-lobar. Caudate. |
| **1** | 4 | 14 | 2 | 0.01 | 2.326348 | Right Cerebrum.Sub-lobar. Caudate. |
| **1** | -12 | 4 | -2 | 0.012 | 2.257129 | Left Cerebrum.Sub-lobar. Lateral Globus Pallidus |
| **Cocaine > Nicotine** | | | | | | |
| **1** | -6.7 | 37.5 | 20.9 | 0 | 3.290527 | Left Cerebrum. Limbic Lobe. Anterior Cingulate. |
| **1** | 0 | 34.7 | 23.3 | 0.001 | 3.090232 | Left Cerebrum. Limbic Lobe. Anterior Cingulate. |
| **2** | 4.7 | 16 | 52.7 | 0.003 | 2.747781 | Right Cerebrum. Frontal Lobe. Superior Frontal Gyrus. |

| | | | | | | |
|---|---|---|---|---|---|---|
| **2** | 3 | 14 | 56 | 0.002 | 2.878162 | Right Cerebrum. Frontal Lobe. Superior Frontal Gyrus. |
| **3** | 22 | 41 | 20 | 0 | 3.290527 | Right Cerebrum. Frontal Lobe. Medial Frontal Gyrus. |
| **3** | 16 | 40 | 24 | 0.006 | 2.512144 | Right Cerebrum. Frontal Lobe. Medial Frontal Gyrus |
| **3** | 7 | 35 | 25 | 0.015 | 2.17009 | Right Cerebrum. Limbic Lobe. Anterior Cingulate. |
| **3** | 14 | 34 | 26 | 0.01 | 2.326348 | Right Cerebrum. Frontal Lobe. Medial Frontal Gyrus. |
| **4** | 2 | 44 | -2 | 0.005 | 2.575829 | Right Cerebrum. Limbic Lobe. Anterior Cingulate. |
| **4** | -6 | 48 | 0 | 0.006 | 2.512144 | Left Cerebrum. Frontal Lobe. Medial Frontal Gyrus. |
| **4** | -4 | 44 | -4 | 0.009 | 2.365618 | Left Cerebrum. Limbic Lobe. Anterior Cingulate. |
| **5** | -46.7 | 24 | 25.3 | 0.015 | 2.17009 | Left Cerebrum. Frontal Lobe. Middle Frontal Gyrus. |
| **6** | -4 | 26 | 2 | 0.003 | 2.747781 | Left Cerebrum. Limbic Lobe. Anterior Cingulate. |
| **6** | -6 | 20 | 4 | 0.011 | 2.290368 | Left Cerebrum.Sub-lobar. Caudate |
| **7** | 38 | 20 | 4 | 0.011 | 2.290368 | Right Cerebrum.Sub-lobar. Insula. |
| **8** | 26 | 2 | 8 | 0.032 | 1.85218 | Right Cerebrum.Sub-lobar. Putamen |
| **Nicotine > Cocaine** | | | | | | |
| **1** | -12 | 8 | 0 | 0.027 | 1.926837 | Left Cerebrum.Sub-lobar. Lateral Globus Pallidus |
| **2** | 30 | 14 | 10 | 0.019 | 2.074855 | Right Cerebrum.Sub-lobar. Claustrum. |
| **3** | -38 | 12 | 2 | 0.008 | 2.408916 | Left Cerebrum.Sub-lobar. Insula. |
| **Alcohol > Nicotine** | | | | | | |
| **1** | -40 | -2 | 52 | 0.027 | 1.926837 | Left Cerebrum. Frontal Lobe. Middle Frontal Gyrus. |
| **1** | -40 | 6 | 52 | 0.039 | 1.76241 | Left Cerebrum. Frontal Lobe. Middle Frontal Gyrus. |
| **2** | 28 | 44 | 18 | 0.019 | 2.074855 | Right Cerebrum. Frontal Lobe. Superior Frontal Gyrus. |
| **3** | -45 | -14 | 47 | 0.014 | 2.197286 | Left Cerebrum. Frontal Lobe. Precentral Gyrus. |
| **Nicotine > Alcohol** | | | | | | |
| **1** | 14 | 16 | 8 | 0.003 | 2.747781 | Right Cerebrum.Sub-lobar. Caudate. |
| **2** | 34 | 16 | -4 | 0.015 | 2.17009 | Right Cerebrum.Sub-lobar. Inferior Frontal Gyrus. |
| **2** | 26 | 22 | 0 | 0.02 | 2.053749 | Right Cerebrum.Sub-lobar. Claustrum. |
| **2** | 30 | 22 | -2 | 0.021 | 2.03352 | Right Cerebrum.Sub-lobar. Insula. |
| **3** | 0 | 14 | 40 | 0.016 | 2.144411 | Left Cerebrum. Limbic Lobe. Cingulate Gyrus. |
| **3** | 2 | 10 | 36 | 0.019 | 2.074855 | Right Cerebrum. Limbic Lobe. Cingulate Gyrus. |

***PostHoc analysis***

**Supplementary Figure 1** shows a side-by-side visual comparison between ALE results due to the studies (labels A1-D1) and task experiments (A2-D2), we found no overlap for nicotine versus alcohol for the task paradigms. **Supplementary Table 2** shows the detail coordinate information for the task-based conjunction. In the chi-square test, we found that the type of experiment may have an influence on the overlaps in alcohol versus cocaine and nicotine versus alcohol, whereas cannabis versus cocaine, cannabis versus nicotine and cocaine versus nicotine are independent of the task paradigms used in the studies shown in **Supplementary Figure 2**. The chi-square values (Yates correction) for $p < 0.05$ are as follows, $\chi^2 = 0.06$, $\chi^2 = 2.6$ and $\chi^2 = 0.03$ respectively.

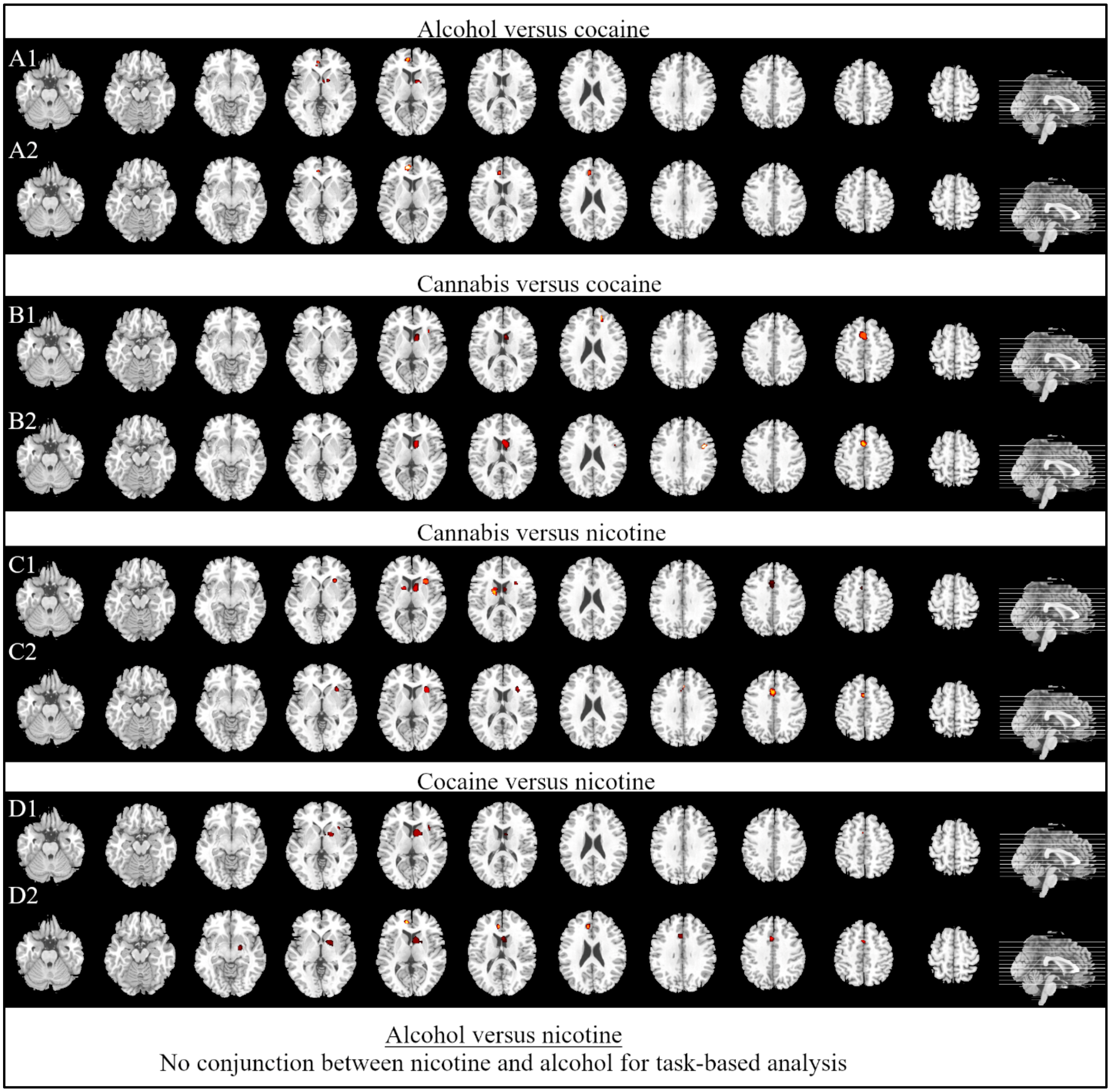

Supplementary Figure 1 Comparing conjunction based on studies and tasks. Each row Figure label shows conjunction based on studies (see Figure 4 in the main paper) and row label 2 shows conjunction based on task respectively

Supplementary Table 2 Detail coordinates forming the conjunction based on tasks. See the main paper Table 3 for studies-based conjunction.

| Cluster # | x | y | z | ALE | Label |
|---|---|---|---|---|---|
| **A2: Alcohol versus cocaine** | | | | | |
| **1** | -8 | 38 | 20 | 0.026185 | Left Cerebrum. Limbic Lobe. Anterior Cingulate. |
| **2** | -8 | 48 | 6 | 0.024981 | Left Cerebrum. Frontal Lobe. Medial Frontal Gyrus. |
| **2** | -6 | 40 | 0 | 0.017812 | Left Cerebrum. Limbic Lobe. Anterior Cingulate. |
| **B2: Cannabis versus cocaine** | | | | | |
| **Cluster #** | x | y | z | | Label |
| **1** | 10 | 8 | 12 | 0.040678 | Right Cerebrum.Sub-lobar. Caudate. |
| **2** | 2 | 10 | 52 | 0.024521 | Right Cerebrum. Frontal Lobe. Superior Frontal Gyrus. |
| **3** | 44 | 4 | 34 | 0.022892 | Right Cerebrum. Frontal Lobe. Precentral Gyrus. |
| **3** | 42 | 2 | 30 | 0.02274 | Right Cerebrum. Frontal Lobe. Precentral Gyrus. |
| **C2: Cannabis versus nicotine** | | | | | |
| **Cluster #** | x | y | z | | Label |
| **1** | 30 | 20 | 8 | 0.030983 | Right Cerebrum.Sub-lobar. Insula. |
| **2** | 0 | 8 | 44 | 0.03443 | Left Cerebrum. Frontal Lobe. Medial Frontal Gyrus. |
| **2** | -4 | 20 | 34 | 0.019912 | Left Cerebrum. Frontal Lobe. Cingulate Gyrus. |
| **3** | 2 | 26 | 30 | 0.016228 | Right Cerebrum. Limbic Lobe. Cingulate Gyrus. |
| **D2: Cocaine versus nicotine** | | | | | |
| **Cluster #** | x | y | z | | Label |
| **1** | 10 | 8 | 8 | 0.02624 | Right Cerebrum.Sub-lobar. Caudate. |
| **1** | 20 | 4 | -2 | 0.025724 | Right Cerebrum.Sub-lobar. Lentiform Nucleus. Putamen |
| **1** | 20 | -6 | -8 | 0.023849 | Right Cerebrum.Sub-lobar. Lentiform Nucleus. Globus Pallidus |
| **2** | -4 | 18 | 32 | 0.023704 | Left Cerebrum. Limbic Lobe. Cingulate Gyrus. |
| **2** | 0 | 10 | 44 | 0.021835 | Left Cerebrum. Frontal Lobe. Medial Frontal Gyrus. |
| **2** | -2 | 12 | 52 | 0.019066 | Left Cerebrum. Frontal Lobe. Superior Frontal Gyrus. |
| **3** | -6 | 38 | 20 | 0.025814 | Left Cerebrum. Limbic Lobe. Anterior Cingulate. |
| **4** | -8 | 46 | 8 | 0.023994 | Left Cerebrum. Frontal Lobe. Medial Frontal Gyrus. |
| **5** | -6 | 38 | 4 | 0.016629 | Left Cerebrum. Limbic Lobe. Anterior Cingulate. |

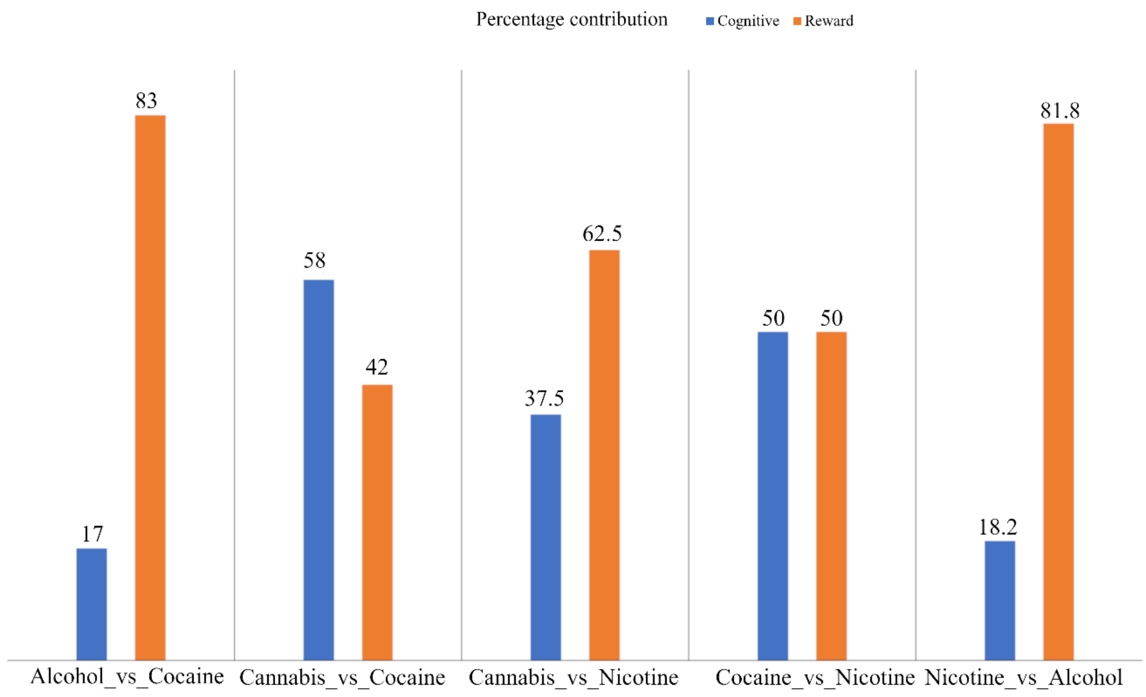

Supplementary Figure 2 Percentage contribution to overlaps based on the task experiment

Supplementary Table 3 Detailed peak cluster information for differences and conjunction between cognitive task and reward task for all substances

| **Cluster #** | **x** | **y** | **z** | **ALE** | **Label** |
|---|---|---|---|---|---|
| **Cognitive task > Reward task** | | | | | |
| **1** | -2 | 10 | 44 | 0.053965 | Left Cerebrum. Frontal Lobe. Medial Frontal Gyrus. |
| **1** | -4 | 16 | 34 | 0.043687 | Left Cerebrum. Limbic Lobe. Cingulate Gyrus. |
| **1** | -2 | -4 | 52 | 0.040893 | Left Cerebrum. Frontal Lobe. Medial Frontal Gyrus. |
| **1** | 0 | 26 | 32 | 0.035746 | Left Cerebrum. Limbic Lobe. Cingulate Gyrus. |
| **1** | 2 | -6 | 42 | 0.035104 | Right Cerebrum. Limbic Lobe. Cingulate Gyrus. |
| **2** | 32 | 18 | 4 | 0.043585 | Right Cerebrum.Sub-lobar. Insula. |
| **2** | 32 | 18 | 8 | 0.041755 | Right Cerebrum.Sub-lobar. Insula. |
| **2** | 38 | 16 | 6 | 0.041452 | Right Cerebrum.Sub-lobar. Insula. |
| **2** | 48 | 16 | 10 | 0.03292 | Right Cerebrum. Frontal Lobe. Inferior Frontal Gyrus. |
| **3** | 44 | 8 | 30 | 0.038719 | Right Cerebrum. Frontal Lobe. Inferior Frontal Gyrus. |
| **3** | 48 | 0 | 38 | 0.034591 | Right Cerebrum. Frontal Lobe. Middle Frontal Gyrus. |
| **4** | -46 | 4 | 34 | 0.052699 | Left Cerebrum. Frontal Lobe. Precentral Gyrus. |
| **5** | 30 | 48 | 14 | 0.047302 | Right Cerebrum. Frontal Lobe. Superior Frontal Gyrus. |
| **6** | -40 | 22 | 0 | 0.036538 | Left Cerebrum. Frontal Lobe. Inferior Frontal Gyrus. |
| **6** | -32 | 20 | 6 | 0.035358 | Left Cerebrum.Sub-lobar. Insula. |
| **7** | -12 | -20 | 8 | 0.038503 | Left Cerebrum.Sub-lobar. Thalamus. |
| **7** | -12 | -8 | 14 | 0.030863 | Left Cerebrum.Sub-lobar. Thalamus. |
| **8** | -2 | 46 | 0 | 0.033395 | Left Cerebrum. Limbic Lobe. Anterior Cingulate. |
| **8** | -8 | 48 | 6 | 0.033241 | Left Cerebrum. Frontal Lobe. Medial Frontal Gyrus. |
| **Reward task > Cognitive task** | | | | | |
| **Cluster #** | x | y | z | ALE | Label |
| **1** | 8 | 4 | 12 | 0.047266 | Right Cerebrum.Sub-lobar. Caudate. |
| **1** | 14 | 6 | 0 | 0.042621 | Right Cerebrum.Sub-lobar. Lateral Globus Pallidus |
| **1** | -10 | -2 | 14 | 0.040866 | Left Cerebrum.Sub-lobar. Caudate. |
| **1** | -8 | 10 | -2 | 0.040428 | Left Cerebrum.Sub-lobar. Caudate. |
| **1** | -10 | 2 | 12 | 0.037655 | Left Cerebrum.Sub-lobar. Caudate |
| **1** | 22 | -6 | -8 | 0.028619 | Right Cerebrum.Sub-lobar. Lateral Globus Pallidus |
| **1** | 14 | -4 | 22 | 0.028066 | Right Cerebrum.Sub-lobar. Caudate. |
| **1** | 24 | -8 | 2 | 0.024022 | Right Cerebrum.Sub-lobar. Lateral Globus Pallidus |
| **1** | 30 | -4 | -2 | 0.022584 | Right Cerebrum.Sub-lobar. Putamen |
| **2** | -6 | 38 | 16 | 0.031412 | Left Cerebrum. Limbic Lobe. Anterior Cingulate. |

| | | | | | |
|---|---|---|---|---|---|
| **2** | 6 | 54 | 12 | 0.030731 | Right Cerebrum. Frontal Lobe. Medial Frontal Gyrus. |
| **2** | -8 | 44 | 8 | 0.030226 | Left Cerebrum. Limbic Lobe. Anterior Cingulate. |
| **2** | 2 | 50 | 24 | 0.03013 | Right Cerebrum. Frontal Lobe. Medial Frontal Gyrus |
| **2** | -6 | 40 | 2 | 0.028048 | Left Cerebrum. Limbic Lobe. Anterior Cingulate. |
| **2** | 18 | 48 | 22 | 0.027496 | Right Cerebrum. Frontal Lobe. Superior Frontal Gyrus. |
| **2** | -2 | 52 | 6 | 0.023568 | Left Cerebrum. Frontal Lobe. Medial Frontal Gyrus. |
| **2** | -6 | 28 | 28 | 0.02356 | Left Cerebrum. Frontal Lobe. Medial Frontal Gyrus. |
| **2** | -8 | 28 | 24 | 0.022809 | Left Cerebrum. Limbic Lobe. Anterior Cingulate. |
| **3** | 0 | 4 | 58 | 0.040162 | Left Cerebrum. Frontal Lobe. Superior Frontal Gyrus. |
| **3** | 0 | 8 | 44 | 0.036585 | Left Cerebrum. Frontal Lobe. Medial Frontal Gyrus. |
| **3** | 2 | 12 | 38 | 0.030312 | Right Cerebrum. Limbic Lobe. Cingulate Gyrus. |
| **3** | -8 | 12 | 56 | 0.024598 | Left Cerebrum. Frontal Lobe. Superior Frontal Gyrus. |
| **3** | -4 | 18 | 34 | 0.02205 | Left Cerebrum. Limbic Lobe. Cingulate Gyrus. |
| **4** | 6 | 24 | 30 | 0.032512 | Right Cerebrum. Limbic Lobe. Cingulate Gyrus. |
| **5** | -2 | -40 | 28 | 0.029772 | Left Cerebrum. Limbic Lobe. Cingulate Gyrus. |
| **5** | -12 | -38 | 30 | 0.021995 | Left Cerebrum. Limbic Lobe. Cingulate Gyrus. |
| **5** | -8 | -30 | 30 | 0.02119 | Left Cerebrum. Limbic Lobe. Cingulate Gyrus. |
| **6** | 42 | 14 | 6 | 0.034024 | Right Cerebrum.Sub-lobar. Insula. |
| **6** | 50 | 2 | 12 | 0.028406 | Right Cerebrum. Frontal Lobe. Precentral Gyrus. |
| **Conjunction** | | | | | |
| **Cluster #** | x | y | z | | Label |
| **1** | 2 | 8 | 53 | | Right Cerebrum. Supp_Motor_Area_R |
| **1** | -4 | 16 | 52 | | Left Cerebrum. Supp_Motor_Area_L |
| **2** | -4 | 12 | 44 | | Left Cerebrum. Limbic Lobe. Cingulate Gyrus. |
| **2** | 4 | 26 | 34 | | Right Cerebrum. Limbic Lobe. Cingulate Gyrus |
| **3** | -4 | 28 | 30 | | Left Cerebrum. Limbic Lobe. Anterior Cingulate |
| **4** | -14 | -4 | 19 | | Left Cerebrum. Sub-lobar. Caudate. |
| **5** | -14 | -4 | 14 | | Left Cerebrum. Thalamus_L |
| **6** | -8 | 50 | 8 | | Left Cerebrum. Frontal_Sup_Medial_L |
| **7** | -80 | 80 | -4 | | Left Cerebrum. Frontal Lobe. Frontal_Mid_L |
